# Intracellular fraction of zona pellucida protein 3 is required for the oocyte to embryo transition in mice

**DOI:** 10.1101/2022.12.23.521410

**Authors:** Steffen Israel, Julia Seyfarth, Thomas Nolte, Hannes C.A. Drexler, Georg Fuellen, Michele Boiani

**Affiliations:** Max Planck Institute for Molecular Biomedicine, Roentgenstrasse 20, 48149 Muenster, Germany; Rostock University Medical Center, Institute for Biostatistics and Informatics in Medicine and Aging Research (IBIMA), Ernst-Heydemann-Strasse 8, 18057 Rostock, Germany

**Keywords:** animal model, embryo development, oocyte, *Trim-away*, zona pellucida

## Abstract

In oocyte biology the zona pellucida has long been known to operate three extracellular functions, namely, encasing the oocytes in ovarian follicles, mediating sperm-oocyte interaction and preventing premature embryo contact with maternal epithelia. The present study uncovers a fourth function that is fundamentally distinct from the other three, being critical for embryonic cell survival. Intriguingly, the three proteins of the mouse zona pellucida (ZP1, ZP2, ZP3) were found abundantly present also inside the embryo four days after fertilization, as shown by mass spectrometry, immunoblotting and immunofluorescence. *Trim-away*-mediated knockdown of ZPs in fertilized oocytes resulted in various grades of embryopathy: knockdown of ZP1 impeded blastocyst expansion, knockdown of ZP2 hampered the 2^nd^ zygotic cleavage, while knockdown of ZP3 hampered the 1^st^ zygotic cleavage - unlike the overexpression. Transcriptome analysis of ZP3-knockdown embryos pointed at defects of cytoplasmic translation in the context of embryonic genome activation. This conclusion was supported by reduced protein synthesis in the ZP3-knockdown and by the lack of cleavage arrest when knockdown was postponed from the 1-cell to the late 2-cell stage. These data place constraints on the long-standing notion that the zona pellucida proteins only operate in the extracellular space. We postulate that such noncanonical localization of ZP3 reflect novel roles in the embryo’s interior during the oocyte-to-embryo transition in mice.

**Graphical abstract:** 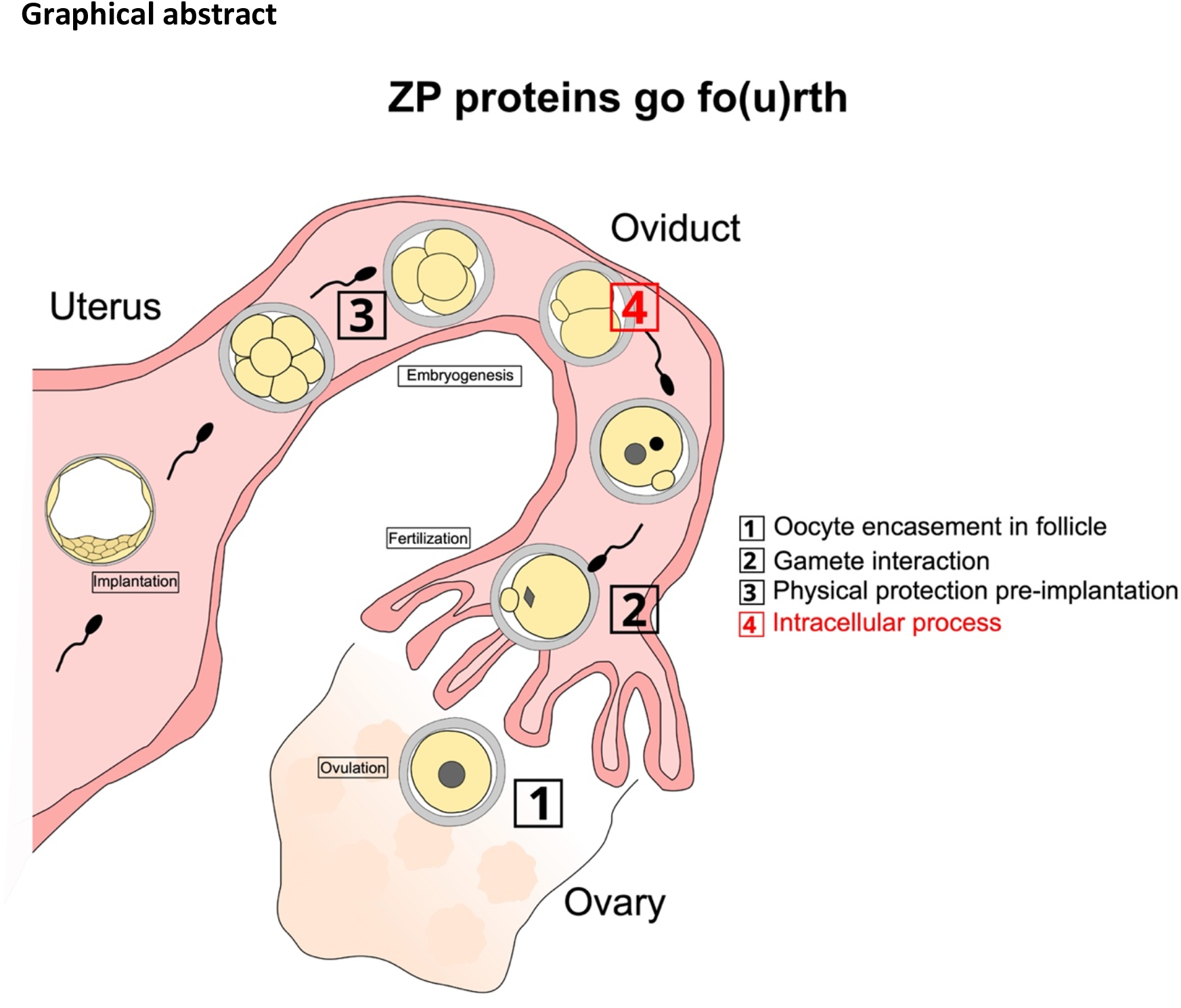

## Introduction

Two of the preconditions for correct embryonic development are that oocytes are protected from physical injury and become fertilized by a single spermatozoon. In mammals these preconditions are secured by the zona pellucida, an extracellular glycoprotein coat encoded by either three or four single-copy genes (e.g. ZP1-3 in mice, ZP1-4 in rabbits and humans, with variations in nomenclature). We are going to use the term ‘zona pellucida’ (‘zona’ for short) to indicate the cytological structure (the extracellular coat), and the acronym ZP to indicate the specific molecular players (genes, transcripts or proteins). During mouse oogenesis the transcription of ZP genes reaches maximum in midsized oocytes (50-60 μm in diameter), before declining to 5% of peak values in ovulated oocytes ^1,2^. ZP genes are not transcribed after fertilization. The immediate translation products are pro-peptides, which are glycosylated and include a signal peptide for routing to the secretory pathway of the oocyte ^1,2^. Owing to these post-translational modifications the molecular masses of each of the three ZP proteins vary, for example: ZP3 weighs 44 kDa as naked polypeptide chain and 83 kDa as glycosylated chain ^2,3^. Once the ZP proteins have been secreted and assembled together, the zona surrounds the embryos until the blastocyst stage, aided by the long half-life of its proteins (>100h; ^4^).

The current model of the zona pellucida builds on the results of ZP mutagenesis studies conducted mostly in mice ^5-8^ but also in rabbits and rats ^9,10^. Based on the mutant phenotypes of ZP2 -/- and ZP3 -/- oocytes, the current model predicates that the zona operates its functions exclusively in the extracellular space. In chronological order these functions are: (1) to encase the oocyte while permitting exchange with the follicular cells via transzonal projections; (2) to mediate species-specific monospermic fertilization, and (3) to hold the blastomeres together while protecting them from premature physical contact with the oviductal epithelium (reviewed in ^11^). In turn, ZP mutations are held responsible for empty follicle syndrome and polyspermic oocyte fertilization ^12^.

A limitation of the above model is that the consequences of the fertilization defects, e.g. polyploidy, precluded analysis of the effects of ZP mutations on embryonic processes. Because the ZP2 -/- and ZP3 -/- oocytes lacked an effective block to polyspermy ^8^, they produced polyploid embryos, which suffer developmental defects regardless of any hypothetical roles played by the ZP proteins after fertilization. To obviate this limitation, investigators fertilized the mutant ZP2 - /- and ZP3 -/- oocytes – which lack the zona - with reduced concentration of sperm *in vitro*, and selected the oocytes that had been fertilized monospermically ^8^. When these diploid zygotes that lacked the zona as a result of the mutation were cultured to blastocyst and transferred to uterus ^8^, birth rates were still severely reduced compared to controls that lacked the zona as a result of manual removal ^8,13^. The reduced competence of ZP -/- oocytes to form viable blastocysts after monospermic fertilization can be explained in two, non-mutually exclusive ways: 1) perduring of oocytes’ defects in the embryo; 2) additional and hitherto uncharacterized functions of ZP2 or ZP3 in the embryonic cytoplasm.

We reasoned that in order to resolve between perduring oocyte defects in the embryo *vs*. new functions of the ZP proteins in the embryo, a postfertilization approach would be beneficial. In plain words, it would be necessary to alter the embryo, with respect to ZP, without altering the precursor oocyte. Since the ZP genes are transcriptionally silenced at the end of oogenesis, a suitable molecular approach is one that hinges on proteins, such as the immunological method of TRIM21-mediated proteasomal degradation, briefly known as ‘*Trim-away*’ ^14^. As we checked how much ZP proteins were present in oocytes, we realized that they were much more abundant than commonly thought, and they also endured throughout preimplantation development. Blastocysts manually freed of the zona still contain almost as much internal ZP proteins as zona-intact blastocysts. Contrary to expectation, knockdown of ZP3 and ZP2 by *Trim-away* in zygotes prevented the 1^st^ and the 2^nd^ cleavage, respectively, while knockdown of ZP1 resulted in stunted blastocyst formation. We then focused on ZP3 *Trim-away*, featuring the earliest and strongest phenotype. Transcriptomic comparison of ZP3-knockdown zygotes *vs*. normal counterparts revealed defects of cytoplasmic translation in the context of embryonic genome activation (EGA). Collectively, this new information supports that ZP proteins are required for developmental competence not only outside but also inside mature oocytes, where the ZPs clearly do more than interacting with each other to form a secreted coat that serves as a scaffold for transzonal projections. We postulate that ZP3 protein is an inner player in the oocyte-to-embryo transition.

## Results

### ZP proteins are abundantly present inside mouse embryos until the blastocyst stage

According to longstanding notion, the zona pellucida exclusively serves its functions outside the oocyte. We were therefore surprised to invariably find, in our previously generated mass spectrometric datasets ^15-19^, conspicuous levels of the proteins ZP1, ZP2 and ZP3 in samples of preimplantation mouse embryos although these had been stripped of their zonae prior to lysis (Supplementary Figure S1). These levels were among the upper percentiles of the proteome abundance distribution ^18^ calculated using the ‘intensity Based Absolute Quantification’ algorithm ^20^, which approximates the molar fraction of the proteins (Figure 1A).

**Figure 1.**
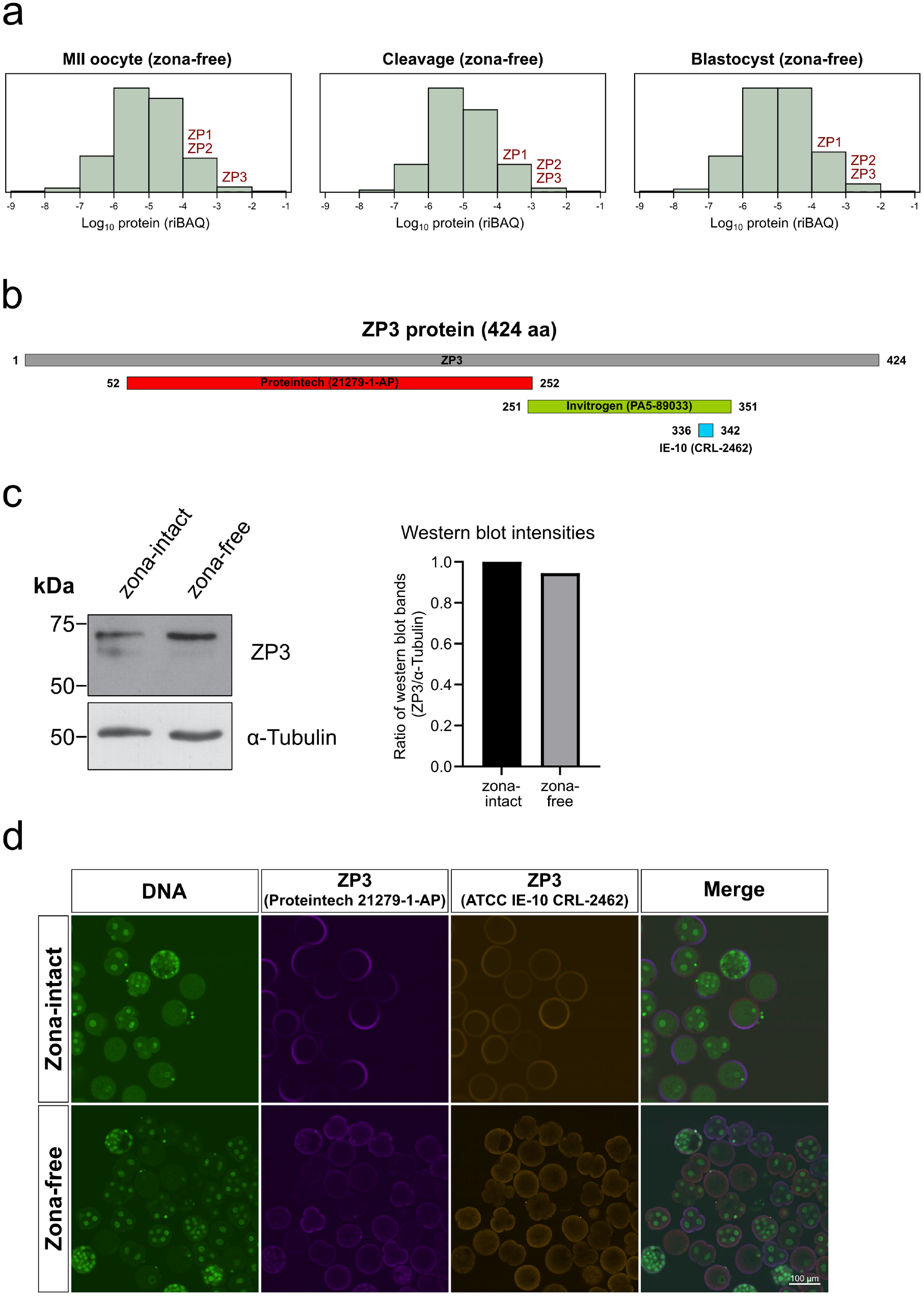
Quantitative and qualitative expression of ZP1, ZP2 and ZP3 during mouse preimplantation development. **(a)**. Genome-wide distribution of protein abundances (riBAQ) in zona-free oocytes, cleavage stages and blastocysts, based on the mass spectrometry dataset PXD012613 ^18^. The proteins ZP1, ZP2 and ZP3 populate, respectively, the 82^nd^, 93^rd^ and 94^th^ percentile of the proteome distribution of oocytes, and the 80^th^, 84^th^ and 84^th^ percentile of the proteome distribution of blastocysts. **(b)**. Three non-redundant antibodies were used to confirm the presence of ZP proteins in the inside of zona-free oocytes and embryos. Shown are the amino acid regions where the antibodies bind to ZP3. **(c)**. Western blot analysis of ZP3 was performed on zona-intact blastocysts (n=400) and zona-free blastocysts (n=544) as well as total ES cell lysate (30 μg) as positive control, using the antibody lnvitrogen PAS-89033. The same blot was processed for ZP3, stripped and reprocessed for α-tubulin as the loading control. Histogram shows the quantified ZP3 bands signals, normalized (zona-intact blastocysts set to 1). **(d)**. Co-immunofluorescence was applied on zona-intact and zona-free embryos of mixed preimplantation stages, using the other two antibodies applied simultaneously (Proteintech 21279-1-AP; ATCC IE-10 CRL-2462). Nuclei (DNA) were stained with YO-PRO-1 and are green fluorescent. The fluorescent signal of the ZP3 protein is not uniformly distributed, neither in the extracellular ring nor in the intracellular fraction.

The unexpected finding of abundant intracellular ZP3 protein throughout preimplantation development was validated with immunoblotting and immunofluorescence using commercially available anti-ZP3 antibodies. A panel of not redundant anti-ZP3 antibodies (binding to different regions of the ZP3 protein) was screened, of which mouse monoclonal ATCC IE-10 CRL-2462, rabbit polyclonal Proteintech 21279-1-AP and Invitrogen PA5-89033 ^18,21-23^ (Figure 1B) were found to work better than others, recognizing a prominent band with ancillary bands of larger size in immunoblots of blastocyst lysates. This composite pattern is expected given the signal peptide and the glycosylation of ZP proteins, as noted in the Introduction. We will be using all three antibodies, although Invitrogen PA5-89033 will be preferred over Proteintech 21279-1-AP in immunoblotting experiments owing to the stronger signal-to-noise ratio; and the rabbit antibodies will be preferred over the mouse antibody in *Trim-away* experiments owing to the higher affinity of TRIM21 for rabbit antibodies ^24^.

In order to validate the strong mass spectrometry signal of ZP3 by an independent method we applied immunoblotting. As before, the zona was completely removed prior to the assay (Supplementary Figure S1). Invitrogen PA5-89033 antibody returned a signal of similar intensity regardless of whether the blastocysts were lysed with or without the zona pellucida (Figure 1C). This was the case also for the other ZP proteins (Supplementary Figure S2). To substantiate this finding while also addressing the subcellular localization of ZP3 we performed co-immunostaining, using the two Proteintech and ATCC antibodies. Co-immunofluorescence stained a ring around the embryos, consistent with common knowledge of the zona as extracellular coat (Figure 1D). However, also the embryos that had been stripped of their zonae pellucida (zona-free) were stained, consistent with the mass spectrometric and immunoblot data. In particular, the intracellular fraction of ZP3 was located peripherally e.g. in the cortical region of the 1-cell stage and in the outer layer (putative trophectoderm) of the blastocyst stage (Supplementary Figure S3A), and the peripheral signal was stronger in some sectors, reminiscent of a polarity (Supplementary Figure S3A). Intracellular staining and subcellular pattern were seen also for the proteins ZP2 and ZP1 (Supplementary Figure S3B).

Together, these findings - enabled by three different methods and distinct antibodies - discount the possibility that the detection of ZP proteins inside the embryos is a false positive. Instead, we hypothesized that there must be a hitherto overlooked intracellular fraction of ZP proteins that persist until the blastocyst stage. Therefore, we became interested in the origin, mechanistic basis and functional meaning of this unusual fraction of ZP proteins found inside the embryos rather than outside.

### Intracellular ZP proteins found in the blastocyst are still those synthesized during oogenesis

Two non-mutually exclusive possibilities are that the intracellular fraction of ZP proteins is a leftover from oogenesis, or that it is produced new in the embryos. The former is based on the principle of parsimony, the latter on the fact that *ZP* genes’ transcripts are present in embryos, as we show hereafter, and that neozona formation was postulated to occur in embryos ^25^. To distinguish between the possibilities, we compared the synthesis of ZP proteins against the abundance of *ZP* transcripts. Although the transcription of *ZP* genes is oocyte-specific and >95% of *ZP3* mRNA is reportedly degraded during ovulation ^2,26^, reanalysis of our previously generated RNAseq data ^18^ revealed that the *ZP* transcripts are in fact not as scarce in embryos as previously assumed. At the blastocyst stage, for example, the *ZP* transcripts populated the median region of the transcriptome distribution (Figure 2A). To resolve if these transcripts are simply present or also translated, we labeled the proteins that are synthesized by the zona-intact zygotes by feeding these with the non-radioactive isotopic amino acids Arg-10 (^13^C_6_H_14_^15^N_4_O_2_) and Lys-8 (^13^C_6_H_14_^15^N_2_O_2_) dissolved in the culture medium. Labeling did not perturb the normal developmental schedule: the zygotes formed blastocyst on day 4 (91% ± 3%, 7 replicates) at a rate similar to control embryos cultured in conventional medium (81% ± 13%, 13 replicates), and the labeled blastocysts also developed to term when transferred surgically to pseudopregnant uteri (216 blastocysts transferred to 27 females, resulting in 14 pregnancies, 59 implantations, 39 fetuses at term). With the safety of this isotopic labeling that preserved developmental potential, we subjected the labeled blastocysts to zona removal followed by proteome analysis. Samples of approx. 500 zona-free blastocysts were collected and subjected to mass spectrometry. In total, 1719 proteins were detected and quantified. A median labeling rate of 83% resulted in 1515 labeled proteins, and the remaining 204 proteins including ZP1, ZP2 and ZP3 were detected as not labeled (Figure 2B; Supplementary Table S1). We next characterized features that distinguish non-labeled proteins, subjecting them to gene and mammalian phenotype ontology analysis using *Enrichr* ^27^. The most prominent terms were “mitochondrial matrix” in the cellular component (GO CC:0005759), “cellular amino acid catabolic process” in the biological process (GO BP:0009063) and “abnormal oocyte morphology” in the mammalian phenotype (MP:0001125). The last term is consistent with the occurrence - among the 204 proteins - of oocyte-specific and maternal-effect genes ^28^ (e.g. *Nlrp2, Nlrp5, Nlrp9, Npm2, Padi6, Tle6, Uhrf1*); we also note their known involvement in mitochondrial function ^29^.

**Figure 2.**
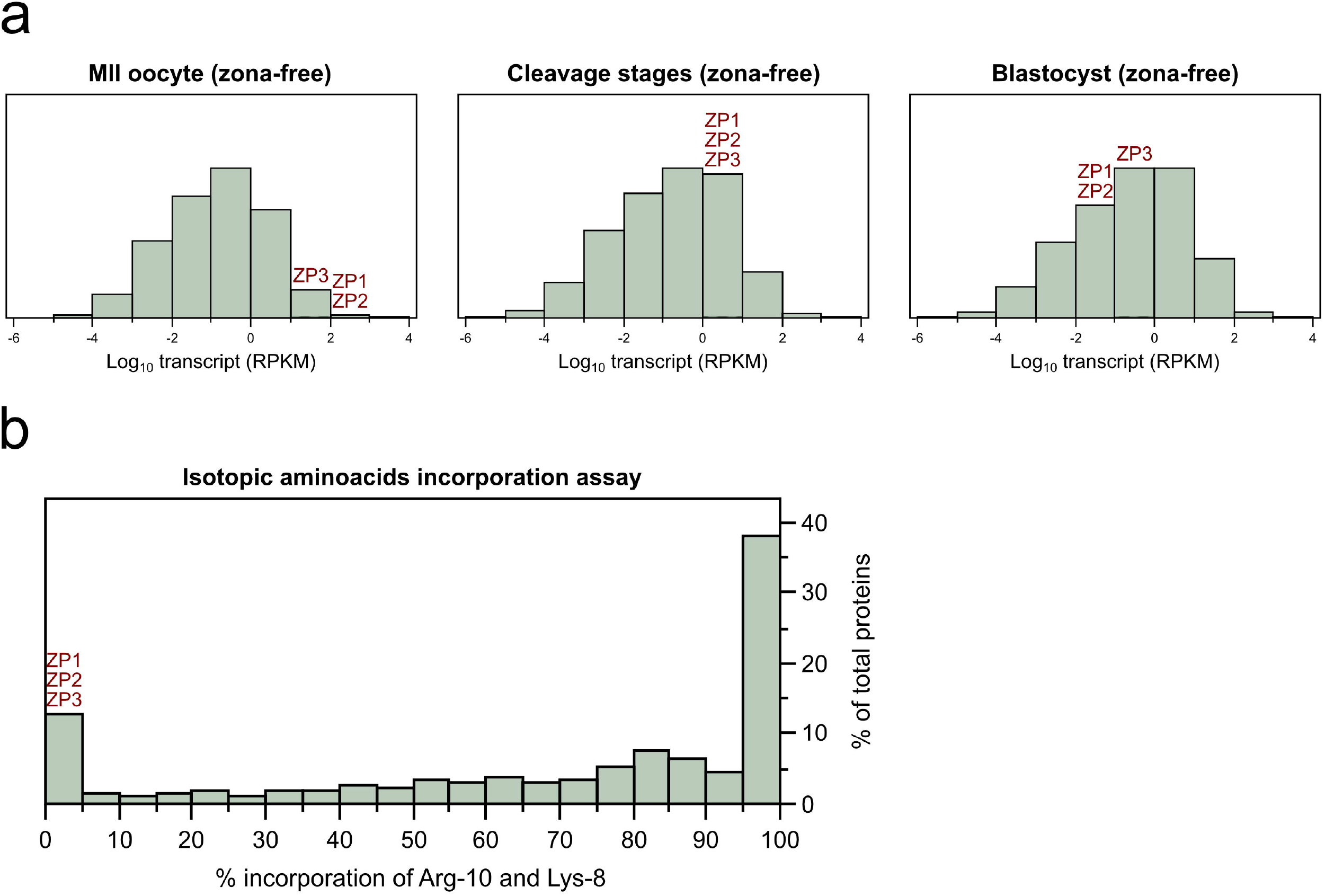
Abundant ZP transcripts are not translated but ZP proteins found in mouse embryos are of maternal origin. **(a)**. Genome-wide distribution of transcript abundances (RPKM) in oocytes, cleavage stages and blastocysts, based on the dataset deposited in the DNA Databank of Japan Sequence Read Archive with the dataset identifier DRA005956 and DRA006335 ^18^. The mRNAs *ZP1, ZP2* and *ZP3* populate, respectively, the 97^th^, 95^th^ and 97^th^ percentile of the transcriptome distribution of oocytes, and the 41 ^st^, 46^th^ and 57^th^ percentile of the transcriptome distribution of blastocysts. **(b)**. When zygotes are cultured four days in the presence of Arg-10 and Lys-8, both amino acids were not incorporated in the ZP proteins found inside the blastocysts, unlike the majority of blastocyst proteins that are labeled by 83% on average (n=1719 proteins; dataset PXD035570). The dataset is available in PRIDE repository with accession number PXD035570 and is provided in simplified form in Supplementary Table S1.

Taken together, these results reveal that although the transcription of ZP genes occurs during oogenesis, the protein products persist during the entire preimplantation phase. This finding elicits considerations about the definition of ‘oocyte-specific’ genes and begs the question as to why embryos hold inside them large amounts of a protein that widespread consensus ascribe with extracellular functions around oocytes.

### An inside job for ZP3: knockdown of ZP3 protein hampered the 1^st^ zygotic cleavage

Given the observation that *ZP* mRNAs are present but not translated in embryos (Figure 2B), it would be futile to test for the developmental relevance of the intracellular ZP proteins by disrupting the DNA loci or depleting the mRNAs. Therefore, we tackled the intracellular fraction directly at the protein level in zona-intact embryos. We followed a dual approach in which ZP3 was inhibited *vs*. knocked down. As pharmacological inhibitors do not exist for ZP3, we microinjected zygotes with defined amounts of either of two anti-ZP3 antibodies, Invitrogen PA5-89033 and Proteintech 21279-1-AP, in purified form (Materials and Methods). Most of the zygotes failed to reach the blastocyst stage when microinjected with Invitrogen PA5-89033 and they even failed to cleave a single time when microinjected with Proteintech 21279-1-AP, being arrested at the pronuclear stage (Table 1; Figure 3A,B) (p = 0.018; Wilcoxon test). The pronucleus-arrested zygotes remained as single cells for four days, without fragmenting nor degenerating (Figure 3A). Developmental arrest was due to the antibody-epitope binding, since blastocyst formation was not impacted when the antibody was heat-denatured ^30^ prior to microinjection (Figure 3A,B). These results lend support to our proposal that ZP3 is needed for hitherto overlooked embryonic processes.

**Table 1.**
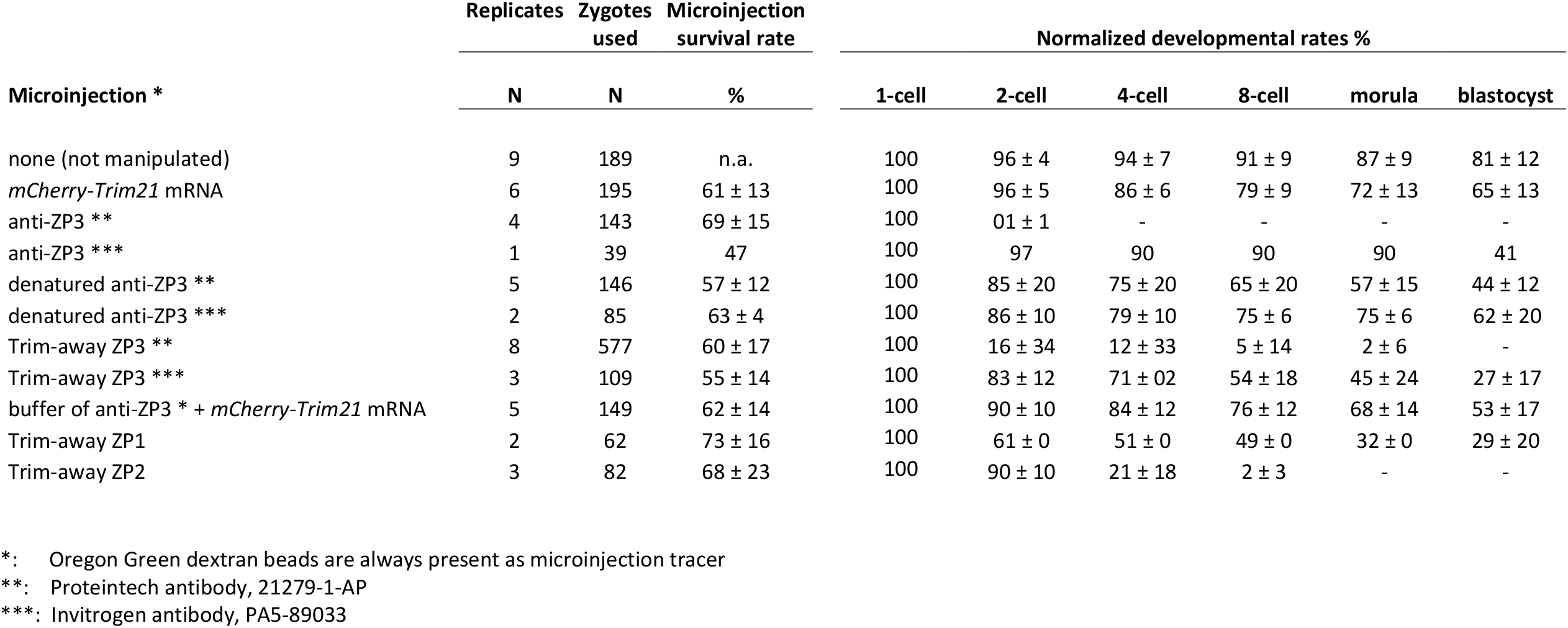
Fertilized mouse oocytes (zygotes) used in the epitope masking and knockdown experiments. Shown are the total numbers of replicates and zygotes manipulated in this study. The developmental rates of the pooled replicates are normalized to the 1-cell stage (zygote). Oregon Green dextran beads were used as tracer in all experiments, to confirm that the microinjection had succeeded technically.

**Figure 3.**
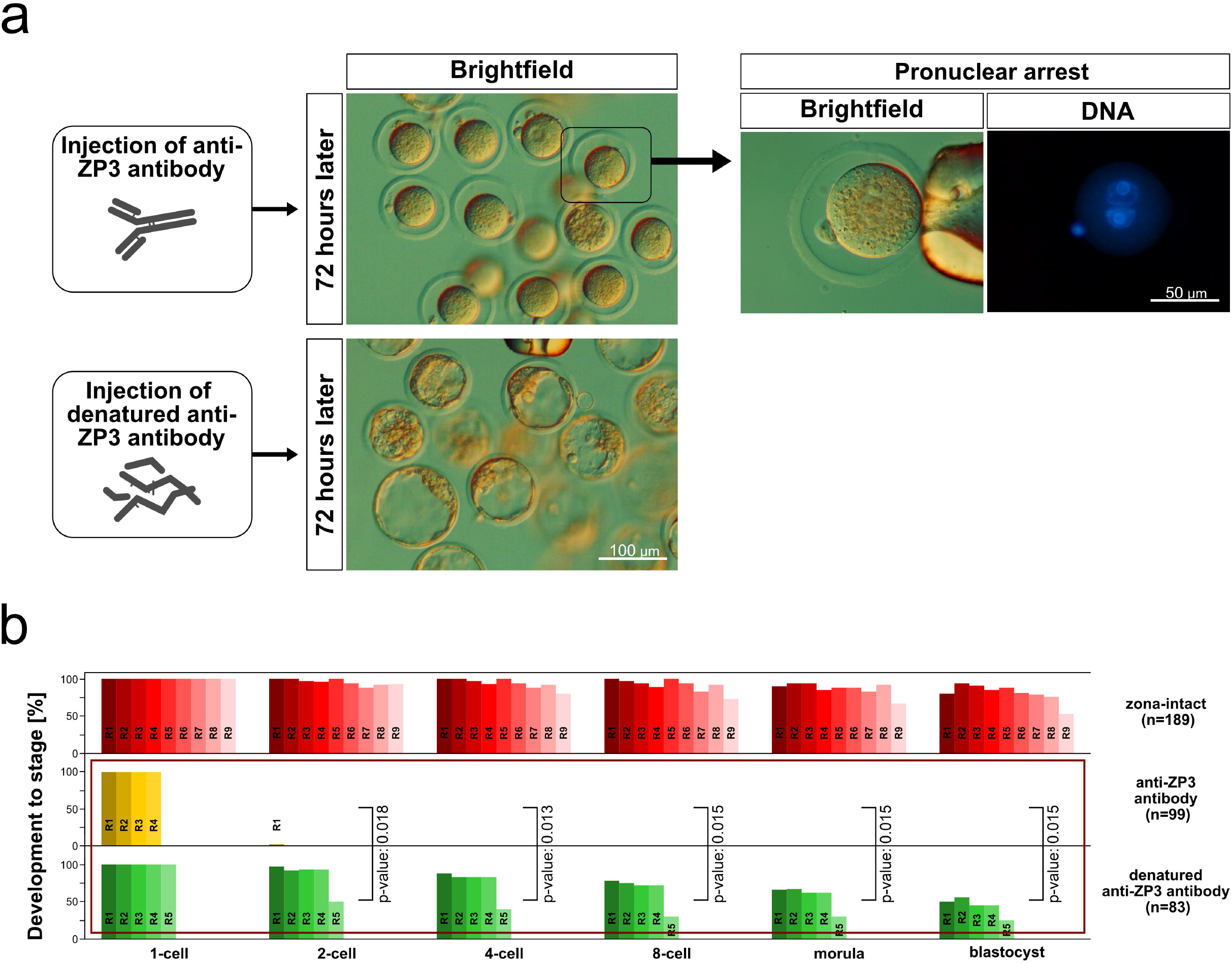
Epitope masking of ZP3 by antibody hampers the 1^st^ zygotic cleavage. **(a)**. Representative images of zygotes that remained arrested at the pronuclear stage when microinjected with anti-ZP3 (Proteintech 21279-1-AP), in contrast to the blastocyst progression of zygotes microinjected with the heat-denatured antibody. Pronuclear-stage arrest was confirmed at higher magnification following Hoechst staining. **(b)**. Developmental rates of the zygotes shown in A, compared to intact zygotes (no microinjection). In each of the three groups the multiple bars next to each other stand for the replicates (anti-ZP3, 4 replicates; denatured anti-ZP3, 5 replicates; non-manipulated i.e. intact, 9 replicates), and the rates of development were normalized to the 1-cell stage (100%). For the replicate of the same group the color is the same, but in different shades. Square brackets between the two series of anti-ZP3 and denatured anti-ZP3 indicate the statistical comparison and its p value (Wilcoxon test). The complete dataset and its breakdown into replicates are provided in Table 1.

Since the epitope masking of ZP3 by the antibody was sufficient to cause 1-cell arrest, we reasoned that ZP3 must support critical cytoplasmic processes. We wondered, therefore, whether the effect would turn out to be even more pronounced when ZP3 is not only masked, but also degraded. To this end, we adopted the method of TRIM21-mediated proteasomal degradation, briefly known as ‘*Trim-away*’ ^14^, which we previously modified for better applicability to large amounts of substrate, such as those characteristically accumulated in oocytes ^15^. The ‘*Trim-away*’ of ZP3 was achieved by microinjection of the same antibody previously used (either Invitrogen PA5-89033 or Proteintech 21279-1-AP), together with the mRNA of the ubiquitin ligase TRIM21 (*mCherry-Trim21* mRNA), thereby triggering the proteasomal commitment of the ternary complex formed between target protein, antibody and mCHERRY-TRIM21. The effect of Proteintech was stronger (Table 1) and this antibody was used preferentially in the subsequent experiments. In our routine setting the reagents are co-injected as a cocktail (Supplementary Figure S4B), which has the advantage to inflict a single microinjection on the zygote. However, this advantage is traded off against the viewing of the consecutive rise and fall of mCHERRY-TRIM21 fluorescence, because mCHERRY-TRIM21 starts to be degraded as soon as the *mCherry-Trim21* mRNA has been translated in sufficient amount. Therefore, we injected the two reagents separately for the sole purpose of visualization: *mCherry-Trim21* mRNA in the zygote, first, followed by ZP3 antibody in one of the two blastomeres originated from the first zygotic cleavage. This allowed us to compare the intensity of mCHERRY-TRIM21 fluorescence in the *Trim-away* blastomere against the other blastomere that contained the same initial amount of mCHERRY-TRIM21: *Trim-away* caused a decline of mCHERRY (Supplementary Figure S4B) which was seen also at the ZP3 protein level (Supplementary Figure S4C). With the comfort of this proof, we went back to the routine microinjection of the cocktail in the zygote. Although the degradation of ZP3 was partial i.e. a knockdown, as revealed by staining of the residual ZP3 with antibody ATCC IE-10 CRL-2462 (Supplementary Figure S4C; Supplementary Table S4), the cellular phenotype was more severe compared to the microinjection of the sole Proteintech antibody (Figure 4A,B; Table 1; p ≤ 0.027; Wilcoxon test): zygotes subjected to *Trim-away* were not only pronucleus-arrested, but also degenerated after 2 days, unlike the zygotes that remained single-celled but alive when receiving the anti-ZP3 antibody alone (Figure 4A). Gene expression analysis will be applied to illuminate the causes (next section, ‘Knockdown of ZP3 interferes with cytoplasmic translation in the context of embryonic genome activation (EGA)’). This effect occurred also when ZP3 was knocked down prior to oocyte activation. We subjected MII oocytes to ‘*Trim-away*’ of ZP3 before activating with Strontium chloride. All treated oocytes remained arrested at the 1-cell stage (N = 0 two-cell embryos/26 oocytes; Proteintech 21279-1-AP). In contrast to the knockdown, the overexpression of ZP3 by way of *mCherry-ZP3* mRNA microinjection was permissive of blastocyst formation (58% of 1-cell embryos, N=85).

**Figure 4.**
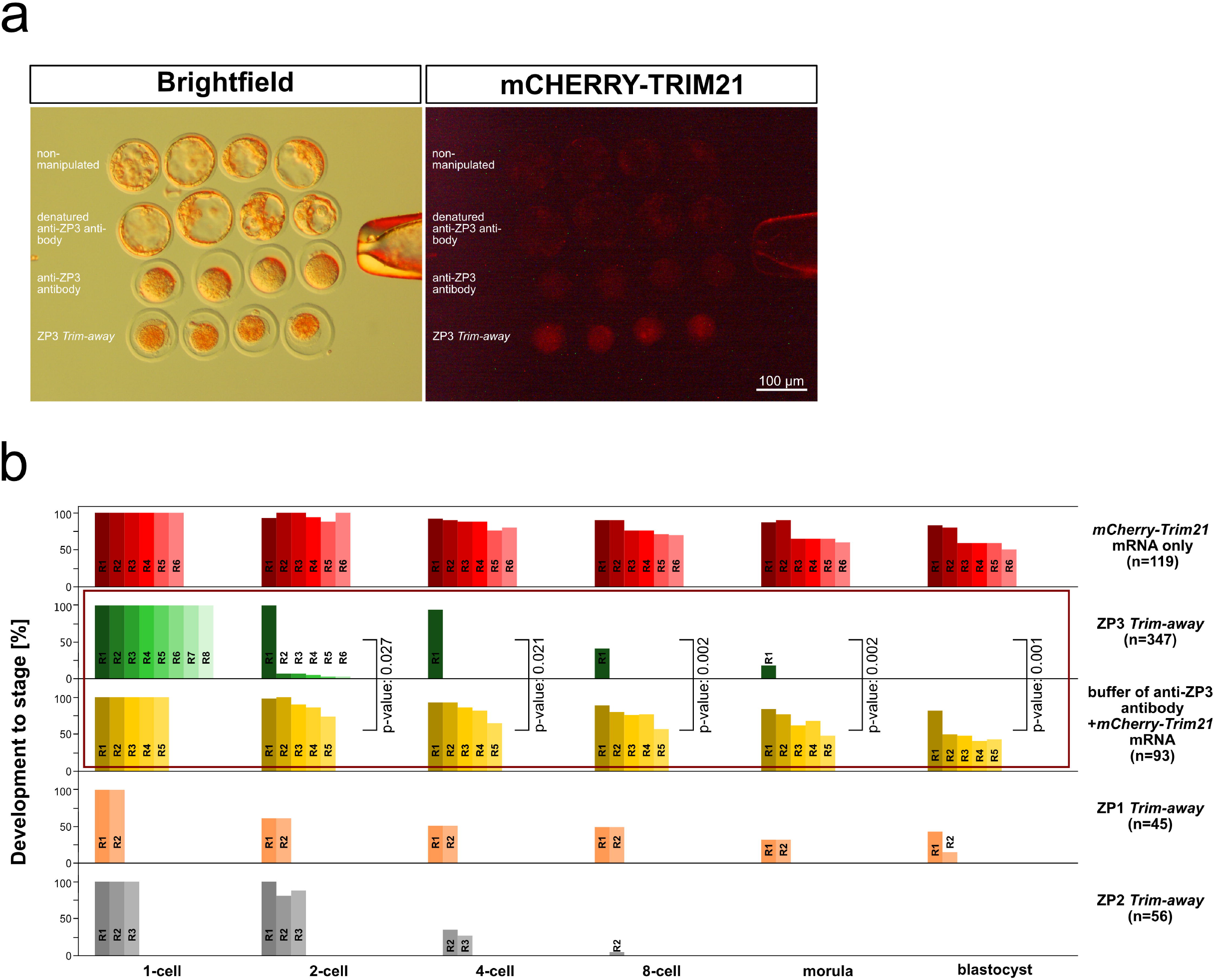
Degradation of ZP3 via *Trim-away* exacerbates the effect of epitope masking by antibody alone. **(a)**. Representative images of zygotes that not only fail to cleave but also degenerate when co-injected with anti-ZP3 antibody (Proteintech 21279-1-AP) and *mCherry-Trim21* mRNA *(Trim-away)*. mCHERRY is the fluorescent tag of TRIM21. **(b)**. Developmental rates of the zygotes subjected to *Trim-away* or its controls e.g. microinjection of *mCherry-Trim21* mRNA only or mRNA with antibody buffer (3^rd^ flow-through of the Amicon filter device). In each of the five series the multiple bars next to each other stand for the replicates *(mCherry-Trim21* mRNA only, 6 replicates; antibody buffer, 5 replicates; ZP3 *Trim-away*, 8 replicates; ZP2 *Trim-away*, 3 replicates; ZP1 *Trim-away*, 2 replicates) and the rates of development were normalized to the 1-cell stage (100%). For the replicates of the same group the color is the same, but in different shades. *Trim-away* of ZP3 hampered the 1^st^ cleavage, *Trim-away* of ZP2 hampered the 2^nd^ cleavage, while *Trim-away* of ZP1 allowed for blastocyst formation but the blastocysts were consistently smaller and less robust than controls. The complete dataset and its breakdown into replicates are provided in Table 1. Square brackets between the two series of ZP3 *Trim-away* and antibody buffer indicate the statistical comparison and its p value (Wilcoxon test).

The pronuclear arrest observed so far is based on the targeting of ZP3. We therefore asked if our findings are due to specific features of this protein, or in other words, whether similar results would manifest also after targeting of ZP1 and ZP2. To this end we applied *Trim-away* to ZP1 and ZP2 (Supplementary Figure S4A, S5). After 4 days, zygotes microinjected with *mCherry-Trim21* mRNA together with anti-ZP2 antibody were found arrested at the 2- and 4-cell stage (0 blastocysts/56 zygotes). Zygotes receiving anti-ZP1 antibody progressed to blastocyst but these were stunted i.e. smaller and less expanded than those from zygotes receiving the sole *mCherry-Trim21* mRNA (18 blastocysts/45 zygotes; Figure 4B; Supplementary Figure S5; Table 1). As in the case of ZP3, these defects were not caused by the microinjection *per se*: zygotes that received the antibody buffer, or the heat-denatured anti-ZP1 or anti-ZP2 antibody progressed efficiently to blastocyst (Table 1).

Taken together, these results reveal that mouse embryos have a hitherto unknown internal requirement for ZP proteins. The ZP proteins found inside are not a leftover from oogenesis, but serve an active role during mouse preimplantation development. Among the three ZP proteins, the requirement of ZP3 is the earliest and strongest, followed by that of ZP2 and then ZP1, in a chronological precession and decreasing severity of phenotype (ZP3 > ZP2 > ZP1). We focused our further molecular investigation of the on the earliest and boldest of the three phenotypes, namely that of ZP3.

### Knockdown of ZP3 interferes with cytoplasmic translation in the context of embryonic genome activation (EGA)

Characterizing the transcriptome is a common way to understand what specific genes are required for. Twenty-four hours past the zygotes’ microinjection with *mCherry-Trim21* mRNA with or without anti-ZP3 antibody we compared the transcriptomes of the two groups of embryos, to illuminate the functions of ZP3 and to identify molecular underpinnings of the pronuclear arrest occurring when ZP3 is knocked down. We also included non-microinjected zygotes, to verify the exogenous supply of *mCherry-Trim21* mRNA in the other two groups, thereby adding up to 8 samples in total (3X *mCherry-Trim21* mRNA only, 4X ZP3 *Trim-away*, 1X non-manipulated; dataset GSE203626; Supplementary Table S2).

Of 17833 gene transcripts detected in total, 11137 were expressed in common in all three groups (Figure 5A; Supplementary Table S2) and were used for further analysis. Embryos that translated mCHERRY-TRIM21 were clustered together with non-microinjected embryos (Figure 5B), documenting that microinjection per se—while being invasive— has a negligible effect on gene expression. Thus, we focused our attention on the significant differences between the *Trim-away* embryos and the embryos that received the sole *Trim21* mRNA. After adjusting for false discovery (p value <0.05 after Benjamini-Hochberg correction), our analysis of the 11137 transcripts identified 198 differently expressed transcripts between ZP3 *Trim-away* embryos and *mCherry-Trim21*-expressing controls. Of the 198 transcripts, 190 also exceeded a 2-fold change (147 down, 43 up; Volcano plot, Figure 5C).

**Figure 5.**
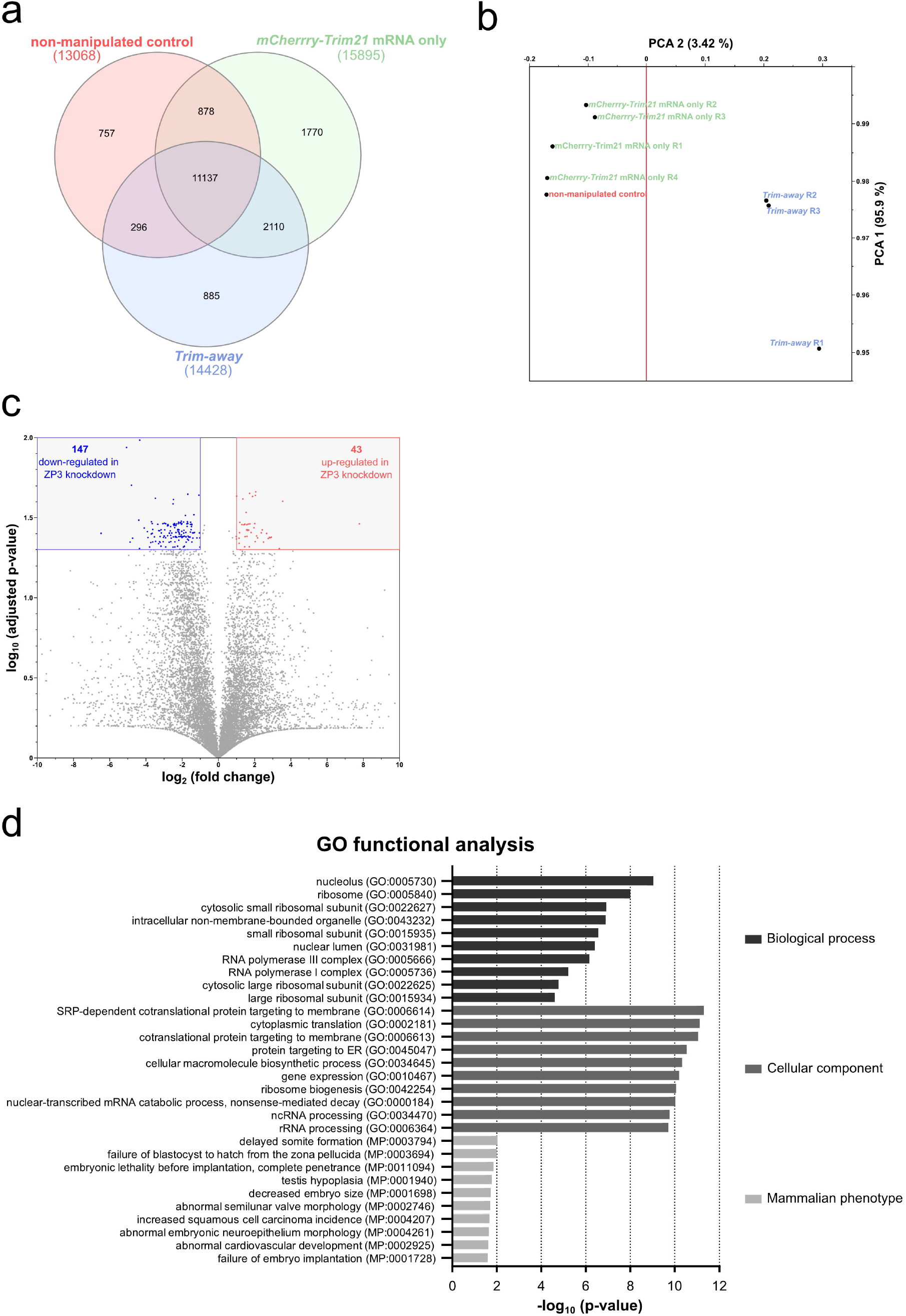
Effects of ZP3 knockdown on the mouse embryonic transcriptome. **(a)**. Zygotes were subjected to transcriptome analysis after microinjection of *mCherry-Trim21* mRNA with or without anti-ZP3 (Proteintech 21279-1-AP), compared to non-injected zygotes. For these three conditions, the Venn diagram describes the overlap in detected transcripts. The RNAseq dataset is available in Gene Expression Omnibus (GSE203626) and is provided in simplified form in Supplementary Table S2. **(b)**. Principal component analysis (PCA) based on the core of 11137 transcripts that are common to all samples. Each point corresponds to one sample *(mCherry-Trim21* mRNA only in triplicate, ZP3 *Trim-away* in quadruplicate, non-manipulated in unicate). The first two principal components (PCs) of the data are represented. The PCA of the transcriptome resolves the ZP3 *Trim-away* and the other groups in the second component. **(c)**. Volcano plot showing the effect of ZP3 *Trim-away* on the embryo transcriptome at the chronological 2-cell stage, compared to *mCherry-Trim21* mRNA alone. The numbers of mRNAs that are under-versus over-expressed (fold change >2, adj.P<0.05, t-test) are shown in the upper corners. **(d)**. The 198 differently expressed mRNAs were subjected to ontology analysis using *Enrichr* ^*27*^, and the top-10 terms (ranked by p-value) were visualized for three ontologies

Since the knockdown of ZP3 caused 1-cell arrest, we asked how the 190 transcripts compare to the catalog of genes whose mutations produce preimplantation lethality in mice, as listed under “MP:0006204” of the Mammalian Phenotype (MP) Ontology. We found seven such genes among the 190 (*Cdca8, Eif6, Kpna7, Kif11, Orc6, Psmc4, Timm23*). This is not a high proportion (4%) yet it suggests that the knockdown of ZP3 compromises the embryos anyway, on multiple levels, and that the pronuclear stage is simply the time when the first process is affected. To illuminate what these processes could be we subjected the set of 190 transcripts to gene and mammalian phenotype ontology analysis using *Enrichr* ^27^. This returned top terms that relate to protein and ribosomal RNA metabolism, such as,SRP-dependent cotranslational protein targeting to membrane’ (GO BP:0006614), ‘cytoplasmic translation’ (GO BP:0002181), ‘nucleolus’ (GO CC:0005730) and,ribosome’ (GO CC:0005840) (Figure 5D), as well as embryo survival, such as ‘failure of blastocyst to hatch from the zona pellucida’ (MP:0003694) (Figure 5D). In regards to ‘cytoplasmic translation’ we recalled that zygotes and early embryos undergo proteome remodeling ^31^ and that interfering with protein synthesis prevents transcriptional activation of the embryonic genome ^32^. Therefore, we measured nascent protein synthesis after incorporation of O-propargyl-puromycin (OPP), as per our established protocol ^16^. The OPP reaction was specific, since it was abolished by preculture with cycloheximide. OPP measurements were performed in 2-cell embryos preloaded with *mCherry-Trim21* mRNA at the 1-cell stage and then microinjected with anti-ZP3 in one blastomere (control group received OGDB in lieu of antibody). This allowed for internal normalization of treatment and control groups. The blastomere that underwent ZP3 knockdown presented lower protein synthesis as captured by the interblastomere difference of fluorescence intensities after Click-chemistry reaction (Figure 6A; Supplementary Table S3). We also recalled that embryos intensify their RNA metabolism to accompany the EGA. Therefore, we searched the 190 transcripts for terms known to be regulated during EGA (Datasets S1-S5 in ^33^), finding 159 of them. Majority of these transcripts (121/159) were mapped to clusters that are upregulated during the minor or major wave of EGA in normal development, while these transcripts were reduced in the pronucleus-arrested zygotes microinjected with ZP3 *Trim-away*. By contrast, the remaining 38 transcripts mapped to the only cluster “consisting of the genes whose expression levels continuously decreased after fertilization and cleavage to the 2-cell stage” ^33^, while these transcripts failed to be downregulated in zygotes microinjected with ZP3 *Trim-away* (Supplementary Figure S6).

**Figure 6.**
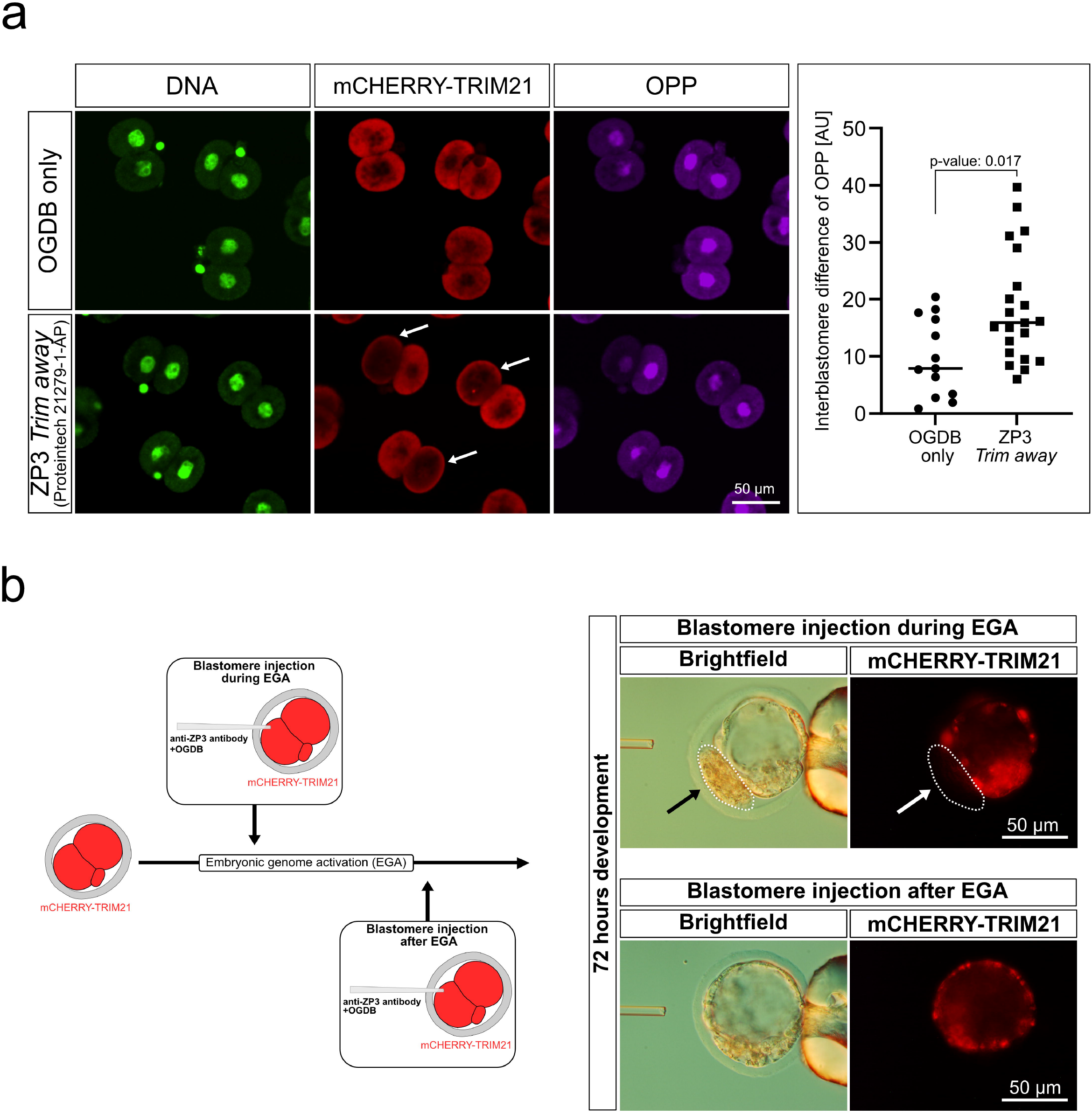
Effects of ZP3 knockdown on cytoplasmic translation and distinct effects of the knockdown depending on the early *vs*. late phase of the 2-cell stage. **(a)**. Zygotes were preloaded with *mCherry-Trim21* mRNA, and after one cleavage they were microinjected with anti-ZP3 (Proteintech 21279-1-AP) or OGDB in one blastomere only, to then be subjected to analysis of protein synthesis using Click-iT ^™^ OPP Alexa Fluor^™^ 647 imaging kit. Note the drop of mCHERRY fluorescence in the blastomere receiving anti-ZP3 (arrows) while the blastomere receiving OGDB retained its fluorescence. OPP fluorescence intensities were measured in both blastomeres of each embryo, and used to calculate absolute interblastomere differences of OPP signal intensity in the *Trim-away vs*. control (OGDB microinjection) groups (p = 0.017; Wilcoxon test). The interblastomere difference is larger in *Trim-away* than in control, because ZP3 knockdown reduces the protein synthesis in one blastomere. The raw measurement data are provided Supplementary Table S3. **(b)**. Using the same design as in (a) the knockdown of ZP3 resulted in developmental arrest when conducted at the early but not at the late 2-cell stage. After 72 h of culture the blastomere that underwent *Trim-away* at the early 2-cell stage produced no cell progeny (arrows and dotted circle), whereas both blastomeres participated in blastocyst formation when *Trim-away* was applied at the late 2-cell stage. Abbreviations: AU, arbitrary units; EGA, embryonic genome activation; OPP, O-propargyl-puromycin.

From the above results it appears that zygotes with knockdown of ZP3 are not apt to manage the RNA and protein dynamics that accompany the oocyte-to-embryo transition. We reasoned that if those changes were the cause of EGA disruption, then *Trim-away* of ZP3 should not matter if conducted at a time when EGA has already occurred. To test this hypothesis, we applied *Trim-away* of ZP3 either early during the 2-cell stage, when EGA is occurring, or late during the 2-cell stage, when EGA has occurred, using the sister blastomeres as reciprocal controls. When *Trim-away* of ZP3 was applied in one blastomere of the early 2-cell embryo it prevented cleavage of the injected blastomere, resulting in a mini-blastocyst formed by the other (untreated) blastomere. By contrast, the blastomere progressed further and participated in blastocyst formation when the same treatment was applied at the late 2-cell stage (Figure 6B).

Collectively, our results corroborate that the ZP3 is required for the oocyte-to-embryo transition in mice. The inhibitory effect on protein synthesis brought about by *Trim-away* of ZP3 is probably the more proximal cause of developmental damage, since treatment with cycloheximide caused 1-cell arrest too. Even if they made it through this hurdle, the zygotes would have a problem shortly after, when the knockdown of ZP3 hampers the major phase of EGA. However, the zygotes are expected to be on the safe side thereafter, as demonstrated by the blastocyst progression of blastomeres subjected to *Trim-away* at the late 2-cell stage, when the bulk of EGA has already occurred.

## Discussion

The significance of the findings presented here is threefold. First, our results expose a hitherto unknown fourth function of the ZP proteins in a mouse model, in addition the traditional functions of 1) encasing the oocytes in ovarian follicles, 2) mediating sperm-oocyte interaction, and 3) protecting the preimplantation embryo from physical injury: as a fourth function there is embryo survival and development. Second, our results document that the newly discovered function of ZP3 is operating intracellularly - an inside job for a protein hitherto believed to operate only outside the cell. Third, our results suggest that the newly discovered function of ZP3 participates in the process of EGA. Collectively, our results sum up to a noncanonical role of ZP3 in oocyte-to embryo transition in mice. The following discussion is centered on the unexpected cytological location and on the novel functional role assigned to ZP3.

As to the unexpected intracellular embryonic location of ZP3 throughout preimplantation development, hints had been there in the past, but they were dismissed as error or trivial case of carryover from oocyte to embryo. Yet *ZP* mRNAs are present throughout preimplantation, in line with suggestions of neozona formation by preimplantation embryos (reviewed in ^25^). Whether neoformation takes place or not was resolved here by performing a live-cell protein experiment of isotopic labeling that has no precedent in preimplantation embryos, although similar work was done on postimplantation stages ^34,35^ and in invertebrates ^36,37^. The finding that the *ZP3* gene product detected in the blastocyst was the oocytic protein that perdured in the embryo, without neoformation, not only challenges the wisdom of calling some genes ‘oocyte-specific’ (clearly not, when by gene product we mean the protein), but also shaped the choice of the method suited to remove the gene product and see what would happen without it. Clearly, we could not use DNA locus disruption or mRNA interference, since the protein of interest is inherited from the mother without embryonic replenishment. As to the functional relevance, ZP3 protein is not only present but also functionally relevant. This knowledge was ‘in the air’ after the report of functional ZP3 found in the germinal vesicles of mouse oocytes ^38^, however, the testing of an embryonic role was more challenging. It was precluded by the confounder of polyspermic fertilization in the mutational studies of ZP3 at a time when the intracytoplasmic sperm injection had yet to become a common tool. In plain words, the ZP3 mutant embryos were crippled with polyploidy and this was the reason why they failed, obscuring possible embryonic roles of ZP3 such as the one uncovered here, namely, a requirement of ZP3 during EGA. This is not the first time a masked phenotype involving ZP3 is uncovered. Upon disruption of the connexin gene *Cx43*, for example, homozygotic mutant mice died shortly after birth ^39^, which precluded their examination at the adult stage. However, when *Cx43* -/- fetal ovary was grafted to wildtype females and allowed to grow, it revealed that mutant oocytes have poorly developed zonae pellucidae ^40^, probably because the transzonal projections that normally contain CX43 ^41^ are needed as scaffolding to assemble the zona.

Given the long-lived maternal deposit of ZP3 revealed by the isotopic labeling experiment, it was clear that the intracellular requirement of ZP3 for embryo survival had to be determined directly in the embryo at the protein level. This was accomplished via methods that either mask the ZP protein (epitope masking) or trigger its proteasomal degradation. Live-cell protein masking by antibodies was introduced in the 1980s to show the effects of disrupting the cytokeratin filament network in mouse embryos ^42^, while the antibody-based proteasomal degradation of proteins is more recent. Known as ‘*Trim-away’*, it was introduced in 2017 and it has been applied by several laboratories ^14,43-47^ including our own ^15,16^ to tackle various proteins of the mammalian oocyte or embryo. In many of those cases the degradation of target protein was partial ^43,44,48,49^, yet a knockdown was sufficient to produce a bold effect, as seen here also with ZP3. In fact, the phenotype of the ZP3 knockdown - pronuclear arrest - was even more severe than that incurred after epitope masking of ZP3: the zygotes not only arrested but also died. It is unlikely that we are dealing with nonspecific effects of a particular match antibody-antigen, since ZP2 and ZP1 subjected to *Trim-away* resulted in embryopathy too.

As if the discovery of ZP3’s fourth function were not surprising enough, it has further implications, first and foremost for the compartmentalization of protein synthesis. It is generally accepted that soluble proteins are synthesized on free ribosomes, whereas secretory proteins – like ZPs - are synthesized on endoplasmic reticulum-bound ribosomes ^50^ and end up facing the luminal side of secretory vesicles. The fact that our microinjection of antibody in the cytoplasm had an effect means that there must be also a subfraction of ZP3 outside of the vesicles, i.e. free in the cytosol. This would be the functionally relevant fraction of ZP3 in the context of the experiments conducted here. It would also explain observations made in other studies, for example, the presence of ZP3 found inside the germinal vesicle of mouse oocytes ^38^.

To shed light on the molecular bases of the newly discovered function i.e. intracellular requirement of ZP3 for mouse embryo survival, we subjected the ZP3-knockdown embryos to transcriptome analysis by RNA sequencing. In order to be sure to measure an effect if present, the analysis was done 24 hours after treatment, although this inevitably means that the ZP3-knockdown zygotes were arrested at the pronuclear stage while the control zygotes (microinjected only with *mCherry-Trim21* mRNA) had divided. This is one limitation of our study. Gene and mammalian phenotype ontology analysis of the perturbed mRNAs returned ‘cytoplasmic translation’ and ‘nucleolus’ as being among the most enriched terms, which is consistent with a known requirement of protein synthesis ^32^ and nucleolar activity ^51^ for the unfolding of the EGA. Indeed, the ZP3-knockdown zygotes incorporated less amino acids in *de novo* protein synthesis, and the detrimental effect of the ZP3 knockdown was confined to the application of *Trim-away* at the time when EGA was in progress. The connection between ZP3 and protein synthesis may raise an eyebrow yet it is consistent with the assignment of ZP3 to the same cluster as ribosomal proteins in the protein atlas ^52^ and in the Harmonizome ^53^. The time confinement, whereby EGA had to be in progress in order for the ZP3 knockdown to exert its effect, has been observed also in case of the orphan nuclear receptor Nr5a2 ^47^. However, while it was logical to find that a nuclear receptor operated in the EGA, the same cannot be said for a protein hitherto considered to be extracellular. Future research will have to look for the interaction partners of ZP3 in the mouse embryo using co-immunoprecipitation, following the example of germinal vesicle-stage oocytes in which ZP3 was shown to interact with nuclear proteins like PTPRK, AIPL1 and DIAPH2 ^38^. This is a second limitation of our study.

To conclude, the finding that ZP proteins are necessary inside the cell for mouse embryo survival changes the game when it comes to naming gene products that are critical to embryos. It also has implications for what defines a gene as oocyte-specific and for medically assisted reproduction. A high level of transcriptional expression in oocytes is often synonymized with specificity or even exclusivity (i.e., the gene is expressed nowhere else). However, the protein may outlive the transcriptional silencing, and be abundant up to the blastocyst stage or even beyond. ZP expression was even reported in testis and in cancer cells ^54,55^. Pending cautious extrapolation to other species, our results expand the spectrum of considerations when advising subfertile couples on the treatment of subfertility via medically assisted human reproduction. When the zona pellucida is abnormal, bypassing the problem of polyspermic fertilization (thin zona pellucida) or fertilization failure (thick or hard zona pellucida) can initially help the couple to produce embryos; but still these might be unable to produce normal blastocysts due to the intracellular intraembryonic requirement of ZPs uncovered here. Depending on which ZP protein is removed or mutated this can lead to earlier or later blockade of embryonic cleavage, no matter if the zona defect was bypassed using intracytoplasmic sperm injection.

## Methods

### Compliance with regulations on research animals

Mice were used for experiments in compliance with the European, German and institutional guidelines as approved by the Landesamt für Natur, Umwelt und Verbraucherschutz (LANUV) of the state of North Rhine Westphalia, Germany (Permit number 81-02.04.2017.A432). Reporting of the animal research findings followed the ARRIVE guidelines ^56^.

### Mouse embryo production

All mice used in this study (N ≈ 250 females B6C3F1, N ≈ 50 males CD1 over a period of ≈ 1 year) were reared in-house at the MPI Münster. They were maintained in individually ventilated type 2 L cages (Ehret), with autoclaved Aspen wood as bedding material and a cardboard tube as enrichment, in groups of 5 females or individually as males. Access to water (acidified to pH 2.5) and food (Teklad 2020SX, Envigo) was *ad libitum*. The animal room was maintained at a controlled temperature of 22 °C, a relative humidity of 55 %, and a 14/10 h light/dark photoperiod (light on at 6:00 a.m.). The hygiene status was monitored every three months, as recommended by Federation of European Laboratory Animal Science Associations (FELASA), and the sentinel mice were found free of pathogens that might have affected the results of this study. Ovulation was induced by intraperitoneal injection of pregnant mare serum gonadotropin (Pregmagon, IDT) and human chorionic gonadotropin (Ovogest, Intergonan). Lean B6C3F1 females aged 8–10 weeks and weighing approx. 25 g were injected i.p. using a 27G needle, at 5 pm, with 10 I.U. eCG and 10 I.U. hCG 48 h apart. The females were mated to CD1 studs aged 3-12 months to produce zygotes, or left unmated to collect MII oocytes. On the next morning the females were killed by cervical dislocation. The cumulus-oocyte complexes were recovered, dissociated in hyaluronidase (cat. no. *151271*, ICN Biomedicals, USA; 50 I.U./mL in HCZB) dissolved in Hepes-buffered Chatot, Ziomek and Bavister medium (HCZB) with bovine serum albumin (BSA) replaced by *polyvinylpyrrolidone (PVP, 40 kDa)* 0.1% w/v. The cumulus-free MII oocytes were cultured in 500 μL of α-MEM medium (Sigma, M4526) supplemented with 0.2% (w/v) BSA, the cumulus-free zygotes in 500 μL of Potassium (K) simplex optimization medium (KSOM) prepared in house as per original recipe ^57^. KSOM contained free aminoacids both essential and non-essential (hence called KSOM(aa)), 0.2% (w/v) BSA and gentamicin (50 I.U./mL), and was used in a four-well Nunc plate without oil overlay, at 37 °C under 6 % CO_2_ in air. To produce parthenogenetic embryos, MII oocytes were activated for 6 h in Ca-free α-MEM medium containing 10 mM SrCl_2_ and 5 μM Latrunculin B (cat. no. 428020, Merck Millipore, Darmstadt, Germany). Following activation, the pronuclear-stage oocytes were washed in KSOM(aa) in three steps of 10 min each to remove the intracellular Latrunculin B accumulated. Parthenotes were cultured in 4-well plates containing 500 μL KSOM(aa) medium at 37 °C (6 % CO_2_).

### Stable isotope labeling of embryonic protein synthesis during preimplantation

Fertilized oocytes were retrieved from oviducts after mating, and cultured in KSOM(aa) medium prepared and used as described before (see ‘Mouse embryo production’), except that: the albumin was replaced with PVP (0.1% w/v) and the two canonical aminoacids L-Arginine (0.3 mM) and L-Lysine (0.2 mM) were replaced with the non-radioactive isotopic forms Arg-10 (^13^C_6_ H_14_ ^15^N_4_O_2_; Cambridge Isotope Laboratories, cat. no. CNLM-539-H-PK) and Lys-8 (^13^C_6_ H_14_ ^15^N_2_O_2_; Cambridge Isotope Laboratories, cat. no. CNLM-291-H-PK). The fertilized oocytes cultured in the presence of Arg-10 and Lys-8 were collected for proteome analysis at the blastocyst stage.

### Transfer of labeled blastocysts to uterus

To confirm that the isotopic labeling preserved the developmental potential of the zygotes, groups of 8 blastocysts were transferred surgically to one uterine horn of pseudopregnant CD1 recipients that had received the copulation plug from vasectomized CD1 males 2 days prior to the embryo transfer. Prior to surgery, CD1 foster mothers were anesthetized with Ketamine (80 mg/kg body weight)/Xylazin (16 mg/kg)/Tramadol (15 mg/kg) in PBS, delivered intraperitoneally. The surgical wounds were sutured with resorbable Marlin violett. Post-surgical pain was alleviated by providing the animals with Tramadol in drinking water (1 mg/mL). Pregnancies were evaluated by C-section just prior to term (embryonic day 18.5).

### Proteome analysis of labeled blastocysts

Following pooling and lysis of the labeled blastocysts in SDS lysis buffer, blastocyst proteins were subjected to rapid acetone precipitation at room temperature in the presence of 20mM NaCl according to the method described by Nickerson & Doucette ^58^. The air-dried pellet was then further processed using the Preomics iST kit according to the manufacturer’s instructions starting with the resuspension of proteins in 20 μl of the kit’s lysis buffer (Preomics, Martinsried, Germany). The digested and purified sample was subsequently dried in an Eppendorf Concentrator and resuspended in Buffer A (0.1% formic acid) for the LC-MS/MS measurement on a Q Exactive HF mass spectrometer online coupled to an EASY nLC 1200 nano-HPLC pump via a Nanospray Flex ion source (Thermo Fisher Scientific). Peptide mixtures were chromatographically separated on a 50cm long fused silica emitter (CoAnn Technologies, MSWil, Netherlands) home-packed with 2.6μm HALO ES-C18 beads (AMT, MSWil, Netherlands) via a linear gradient from 3% Buffer B (80% acetonitrile, 0.1% formic acid) to 35% B within 220 min, before being ramped to 60% and 98% B within 20 min and 3 min, respectively (flow rate 300 nl/min). Data were recorded using a top 17 data-dependent method (scan range 300-1750 m/z; MS1 resolution 60,000; AGC target 3e6; maximum IT 100 ms; MS2 resolution 15,000; AGC target 1e5; maximum IT 50 ms; normalized collision energy = 27 V; dynamic exclusion enabled for 20 s). Raw data were processed for identification and quantification by MaxQuant Software (version 2.0.3.0) with the ‘iBAQ’ option enabled and the ‘requantify’ option disabled. MaxQuant provides intensities for heavy and light labelled peptides and proteins separately and this also pertains to the iBAQ values. The search for identification was performed against the UniProt mouse database (version from 04/2019) concatenated with reversed sequence versions of all entries and supplemented with common lab contaminants. Parameters defined for the search were trypsin as the digesting enzyme, allowing two missed cleavages, a minimum length of six amino acids, carbamidomethylation at cysteine residues as a fixed modification, oxidation at methionine, and protein N-terminal acetylation as variable modifications. The maximum mass deviation allowed was 20 ppm for the MS and 0.5 Da for the MS/MS scans. Protein groups were regarded as identified with a false discovery rate (FDR) set to 1 % for all peptide and protein identifications; at least one unique peptide was required for each protein analyzed further. The molar fractional content of each protein P in a sample (relative iBAQ intensities = riBAQ_P_) were determined for the light and heavy labeled proteoforms independently according to Shin et al. ^20^, as follows:

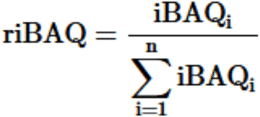

### Immunofluorescence analysis of ZP1, ZP2 and ZP3 expression in preimplantation embryos

Oocytes or embryos were analyzed by performing an immunostaining followed by confocal microscopy imaging, as per our routine protocol. Briefly, embryos were fixed with 3.7 % formaldehyde in 1x PBS for 20 min at room temperature, permeabilized with 0.1% Triton X-100 in 1x PBS for 20 min at room temperature, and then blocked with 2 % BSA, 2 % glycine, 5 % donkey serum, 0.1 % Tween20 in 1x PBS overnight at 4 °C. When applicable, the zona was removed by bathing the oocytes or embryos in acidic Tyrode solution prewarmed at 30 °C (Sigma, T1788). The following primary antibodies were applied to the specimens for 2 h at RT: anti-ZP1 polyclonal IgG raised in rabbit (Invitrogen PA5-101973), anti-ZP2 polyclonal IgG raised in rabbit (ABclonal, A10126), anti-ZP3 polyclonal IgG raised in rabbit (Proteintech 21279-1-AP), anti-ZP3 monoclonal IgG raised in mouse (ATCC IE-10 CRL-2462; 0.5 mg/ml), in dilutions of 1:100, respectively. An appropriate Alexa Fluor-tagged secondary antibody (Invitrogen) was matched to the primary and incubated for 1-2 h at room temperature. DNA counterstaining was performed with YO-PRO-1 (1 μM). For imaging, embryos were placed in 5 μl drops of PBS on a 50-mm thin-bottom plastic dish (Greiner Bio-One, Lumox hydrophilic dish; Frickenhausen, Germany) and overlaid with mineral oil (M8410 Sigma). Images were captured using a 20x CFI Plan Apochromat VC objective on an inverted motorized Nikon TiE2000 microscope fitted with an Andor Dragonfly 502 spinning disc confocal unit Scanning System. Optical sections per embryo were captured using a high resolution (2048 × 2048 pixel) sCMOS camera. Maximum projections were analyzed with Fiji ^59^.

### Immunoblotting analysis of ZP1, ZP2 and ZP3 expression in preimplantation embryos

Oocytes or embryos were centrifuged in protein-free HCZB medium at 39 G for 20 min to form a tiny pellet (Supplementary Figure S1). The supernatant was carefully aspirated using a mouth-operated micropipette, and replaced by RIPA buffer containing protease inhibitors. The resultant lysates were mixed with 6x Laemmli sample buffer and boiled for 5 min at 99 °C. These samples were loaded on a 12 % separation gel and blotted onto a PVDF membrane. The membrane was blocked for at least 3 h and incubated (3 % nonfat dry milk in 0.1 % PBS-Tween20) with primary antibodies overnight at 4 °C. The antibodies against the proteins of interest (ZP3, Anti-ZP3: Invitrogen Catalog # PA5-89033; ZP2, Anti-ZP2: ABclonal Catalog # A10126; ZP1, Anti-ZP1: Invitrogen Catalog # PA5-101973) were applied at a dilution factor of 1:2000 (ZP3), 1:1000 (ZP2) and 1:80 (ZP1). Signal intensities were standardized on α-Tubulin (1:5000, Merck, Cat. no.: T6199). After 3X washing in 0.1 % PBS-Tween20, the blot was incubated with horseradish peroxidase (HRP)-coupled secondary antibody at RT for 1 h. The membrane was washed and then developed with chemiluminescent HRP substrate solution. The chemiluminescent signal was detected using the AGFA Curix 60.

### Epitope masking and proteasomal degradation of ZP proteins

In order to mask the protein of interest, zygotes were microinjected with ZP3 antibody (Proteintech 21279-1-AP) tohether with dextran beads fluorescently labeled with Oregon Green (OGDB; 70 kDa; ThermoFisher cat. no. D7173) at 16 h post-hCG. In order to also degrade the protein, zygotes, early 2-cell embryos and late 2-cell embryos were microinjected with a mixture of *mCherry-mTrim21* mRNA, OGDB and ZP1 (Invitrogen PA5-101973), ZP2 (ABclonal A10126) or ZP3 antibody (Proteintech 21279-1-AP; Invitrogen PA5-89033; ATCC IE-10 CRL-2462) at approximately 16 h, 40 h and 48 h post-hCG, respectively. Concentrations in the mixture were 0.15 mg/mL mRNA, 0.017 mg/mL OGDB and 1 mg/mL antibody, respectively, in MilliQ water. The mRNA was purified with Quick-RNA MicroPrep (Zymo Research, cat. no.: R1051) and preserved in MilliQ water at -80 °C. The antibody was washed three times and concentrated at 4 °C using Amicon Ultra-0.5 100 KDa centrifugal filter devices (Merck Millipore, cat. no. UFC100), which remove salts and preservatives (e.g. sodium azide) and stabilizers (e.g. bovine serum albumin), replacing most of the antibody buffer with water. This buffer was collected as flow-through of the Amicon device and set aside for later use as control. Microinjection of the mRNA-antibody-OGDB mixture was conducted on the stage of a Nikon TE2000U microscope fitted with a piezo drill (PrimeTech), using a blunt-end glass needle (inner diameter 6-7 microns, outer diameter 8-9 microns) filled with 2-3 microliters mercury at the tip. Volumes were pressure-injected into the zygote or blastomere using a Gilmont GS-1200 micrometer syringe operated manually. During the microinjection, cells were kept in a 200-300 microliters drop of HCZB medium on a glass-bottomed (Nomarski optics) dish at a room temperature of 28 °C. After microinjection, zygotes and 2-cell embryos were allowed to recover in the drop for 5–10 min, before returning them to KSOM(aa) medium.

### Synthesis of mRNA for microinjection

For *Trim-away*, an *mCherry-Trim21* expression construct built on plasmid pGEMHE-mCherry-mTrim21 was obtained from Melina Schuh (Addgene plasmid # 105522). For ZP3 overexpression, the coding sequence of *ZP3* (NM_011776.1) was substituted for that of *Trim21* in the plasmid pGEMHE-mCherry. The construct is apt to support strong transcription given the presence of Kozak 5’, *Xenopus* 5’ Globus and *Xenopus* 3’ UTR sequences. For *in vitro* transcription, plasmids were linearized with SwaI (ThermoFisher, cat. no. FD1244). Capped mRNA was synthesized with T7 polymerase (Ambion mMessage mMachine T7 kit) according to manufacturer’s instructions.

### Transcriptome analysis of embryos sampled 24 h after ZP3 knockdown

To assess the consequences of ZP3 inactivation on embryonic gene expression at the chronological 2-cell stage we generated a transcriptomic dataset. Samples were comprised of 10 embryos in each of two groups: group 1 injected with *mCherry-Trim21* mRNA and OGDB and solvent of the ZP3 Proteintech antibody (triplicate), and group 2 injected with *mCherry-Trim21* mRNA, OGDB and anti-ZP3 Proteintech antibody (quadruplicate). One additional sample of embryos without any treatment was also included as an outgroup comparison. Total RNA was extracted and purified using Quick-RNA MicroPrep (Zymo). The library preparation of the total RNA was performed with the NEBNext® Single Cell/Low Input RNA Library Prep Kit for Illumina (NEB #E6420S/L). Single read sequencing with a read length of 72 bp was performed on NextSeq ® 2000 System using the corresponding NextSeq2000 P3 Reagent Kit (Illumina). Total RNA integrity and quality of the library were assessed using a TapeStation4200 (Agilent). On average the libraries contained 18.4 +/- 3.6 million 72-base-single-end reads. Using a molecular barcode, the samples were demultiplexed and converted to fastq data using bcl2fastq v3.8.4 (Illumina) quality controlled (FastQC ^60^; 08-01-19: Version 0.11.9 released). Trimmomatic was used for adapter trimming and read filtering ^61^. The resulting reads were aligned to the reference genome (*Mus musculus* Ensembl GRCm38) using Hisat2 ^62^. The aligned reads were sorted using samtools ^63^. The sorted and aligned reads were counted into genes using htsec-counts ^64^. Then these counts were normalized for library size and transcript length using the RPKM method (Reads Per Kilobase of transcript per Million mapped reads) to normalize for sequencing depth and gene length. RPKM values for Ensembl gene identifiers corresponding to the same gene symbol were averaged, and the values associated with each gene symbol were averaged across replicates.

### Functional enrichment analysis of differently expressed mRNAs

Functional enrichment analysis was performed with *Enrichr* at https://maayanlab.cloud/Enrichr/ ^27^. Apart from the Gene Ontology (GO) of biological processes (BP) and cellular components (CC), we selected mammalian phenotype (MP) ontology, which builds on the Mouse Genome Informatics database and is, therefore, well-suited to examine genes relevant to the mouse and its developmental biology. Terms with a FDR ≤ 0.01 were considered enriched.

### Detection of *de novo* protein synthesis

O-propargyl-puromycin (OPP) was added to the culture medium KSOM(aa) at a final concentration of 20 μM, and the embryos were cultured for 3h. For inhibition of protein synthesis (negative control) embryos were precultured in 50 μg/ml cycloheximide as described ^17^, before switching to KSOM(aa) with OPP. The incorporation OPP was revealed using a Click-iT™ OPP Alexa Fluor™ 647 imaging kit (cat. no. C10458, Invitrogen, Thermo Fisher Scientific, Karlsruhe, Germany), according to the manufacturer’s protocol. Following this procedure, all embryos were fixed with 3.7 % formaldehyde for 15 min, followed by a 0.5 % Triton X-100 permeabilization step for 15 min at room temperature and then incubated with the Click-iT reaction cocktail for 30 min, protected from light. Images were taken with an Andor Dragonfly spinning disc confocal unit Scanning System as described.

### Statistical analysis of developmental rates, images and RNA sequencing results

Developmental rates of embryos and image intensities of immunoflorescence were analyzed non-parametrically by Wilcoxon test using the statistical program JMP Pro v.16 (SAS). RNAseq data analysis was performed in-house using the output of the NextSeq ®2000 System, exported in Microsoft Excel format and imported in JMP Pro v. 16. Differently expressed transcripts were identified using the t test and the resulting P-values were corrected utilizing Benjamini-Hochberg’s method (false discovery rate < 0.05).

## Acknowledgements

The authors would like to express gratitude for scientific environment and infrastructural support to the Max Planck Institute for Molecular Biomedicine. As part of infrastructural support, we experienced outstanding support from the mouse housing facility, ensuring a dependable supply of mice needed to collect oocytes and embryos. The authors thank Annalen Nolte and Line Lüken for help with the LC-MS/MS measurements. The RNAseq analysis was outsourced to the Core Genomic Facility of the Faculty of Medicine of the University of Muenster (Anika Witten, Andreas Huge). A heartfelt thank you to Francesca Duncan for reading the manuscript. This work was supported by the Deutsche Forschungsgemeinschaft (grant DFG BO-2540/8-1 to M.B. and grant FU 583/7-1 to G.F.) and was in part presented orally to the ESHRE meeting 2022 ^65^.

## Author contributions

S.I. and M.B. conceived and co-designed the study. S.I. synthesized the mRNAs, and purified the mRNAs and proteins for microinjection, together with J.S. and T.N. T.N. did the Western blots. H.D. performed the mass spectrometry analysis. M.B. performed the microinjections, embryo culture, embryo transfers, analyzed the results including the statistical analysis with help from S.I. S.I. drew the figures. M.B. wrote the manuscript together with G.F. All authors approved the manuscript.

## Data availability statement

All large-scale data generated during this study are provided in this article as deposited datasets and/or summary tables (Supplementary Tables S1-2). The RNA-seq dataset of ZP3 *Trim-away* embryos at the chronological 2-cell stage has been deposited in the Gene Expression Omnibus of NCBI with accession number GSE203626. To review GEO accession GSE203626 go to https://www.ncbi.nlm.nih.gov/geo/query/acc.cgi?acc=GSE203626 and enter token mhoruysoplczdgz into the box. The mass spectrometry dataset of blastocysts obtained after stable isotope labeling with Lys-8 (^13^C_6_ H_14_ ^15^N_2_O_2_) and Arg-10 (^13^C_6_ H_14_ ^15^N_4_O_2_) in vitro has been deposited in the PRIDE repository with accession number PXD035570 (Username; Password: gOGiJIXA). For convenience, the large-scale data are summarized in the Supplementary Tables S1 and S2.

## Additional information

### Ethics declaration for human experiments and consent for publication

Not applicable.

### Ethics declaration for animal experiments

Mice were used for experiments according to the license issued by the Landesamt für Natur, Umwelt und Verbraucherschutz of the State of North Rhine-Westphalia, Germany (license number LANUV 81-02.04.2017.A432), in accordance with the procedures laid down in the European Directive 2010/63/EU. We observed the ARRIVE guidelines ^56^ to the extent applicable.

