## Supplementary Figures S1-S6 for "Intracellular fraction of zona pellucida protein 3 is required for the oocyte to embryo transition in mice"

**Running title:** An inside job for ZP3

Steffen Israel [1], Julia Seyfarth [1], Thomas Nolte [1], Hannes C.A. Drexler [1], Georg Fuellen [2] and Michele Boiani [1,*]

[1] Max Planck Institute for Molecular Biomedicine, Roentgenstrasse 20, 48149 Muenster, Germany

[2] Rostock University Medical Center, Institute for Biostatistics and Informatics in Medicine and Aging Research (IBIMA), Ernst-Heydemann-Strasse 8, 18057 Rostock, Germany

**Supplementary figures**

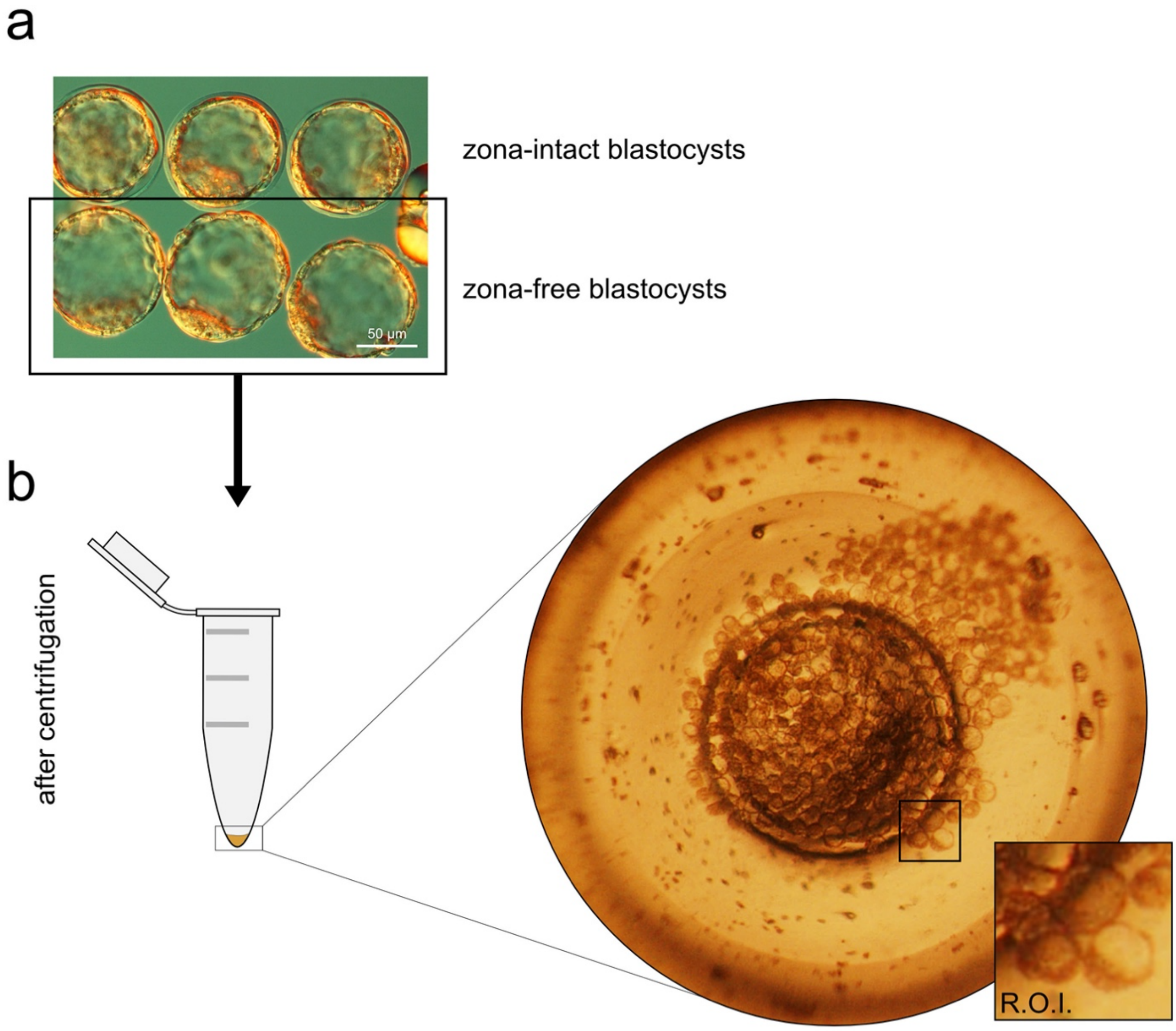


**Supplementary Figure S1. Zona-free sample preparation. (a).** Zona-free blastocysts prior to lysis for mass spectrometry or Western blotting. **(b).** Stereomicroscopic view of a pellet of zona-free blastocysts (n ≈ 200) collected on the bottom of a tube after centrifugation, prior to further processing for mass spectrometry or Western blotting. R.O.I. shows the blastocysts at higher magnification inside the tube. Zona pellucida removal was performed using acidic Tyrode solution. R.O.I., region of interest.

a

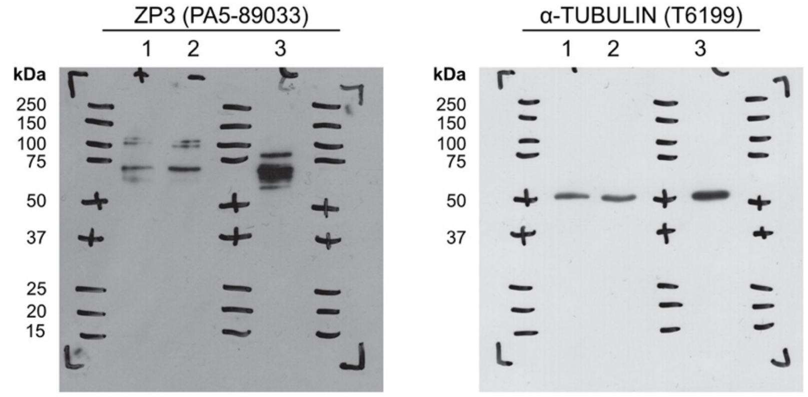


Key
1. #400 non-manipulated blastocysts zona intact
2. #544 non-manipulated blastocysts zona-free
3. 30 µg ES cell protein lysate

b

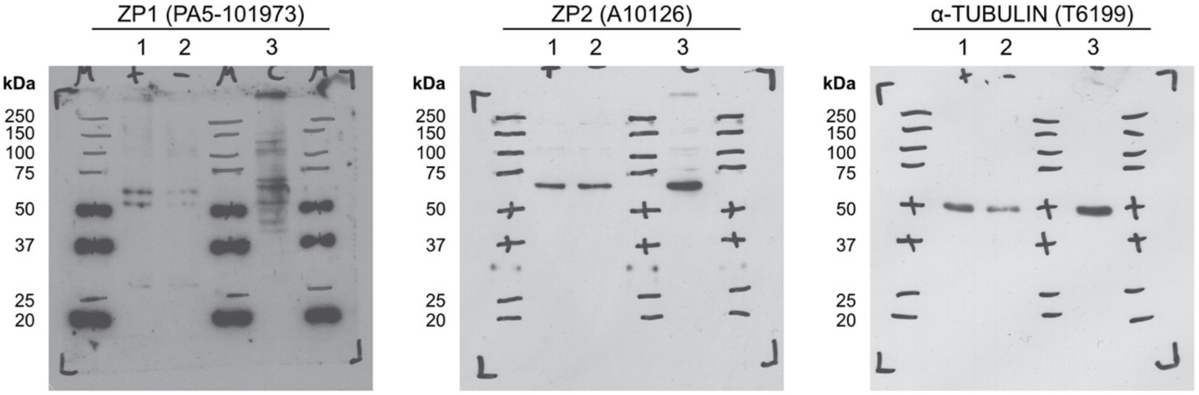


Key
1. #503 non-manipulated blastocysts zona intact
2. #500 non-manipulated blastocysts zona-free
3. 30 µg ES cell protein lysate

**Supplementary Figure S2. Uncropped western blots of ZP1-3. (a).** Uncropped western blots related to ZP3 in Figure 1C. **(b).** Uncropped western blots showing similar pattern also for ZP2 and ZP1, namely: the zona-free blastocysts contain almost the same amount of ZP protein as the zona-intact blastocysts, indicating that the intracellular fraction of ZPs is larger than the extracellular fraction. The same blot in (a) and the same blot in (b) were stripped and reprocessed for α-tubulin as the loading control. For reasons unknown the anti-ZP1 antibody binds also the molecular weight ladder.

a

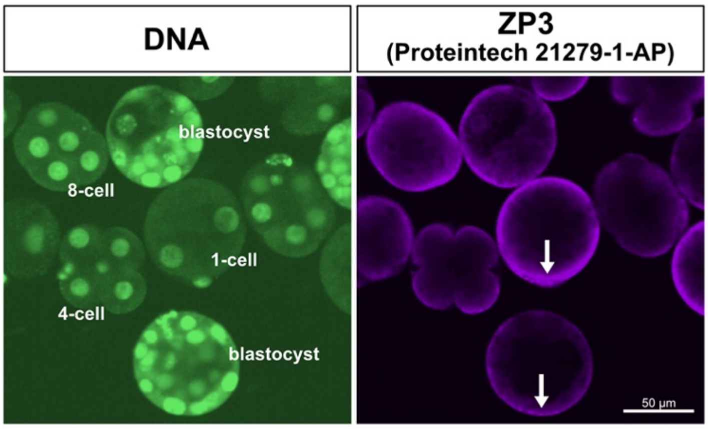


b

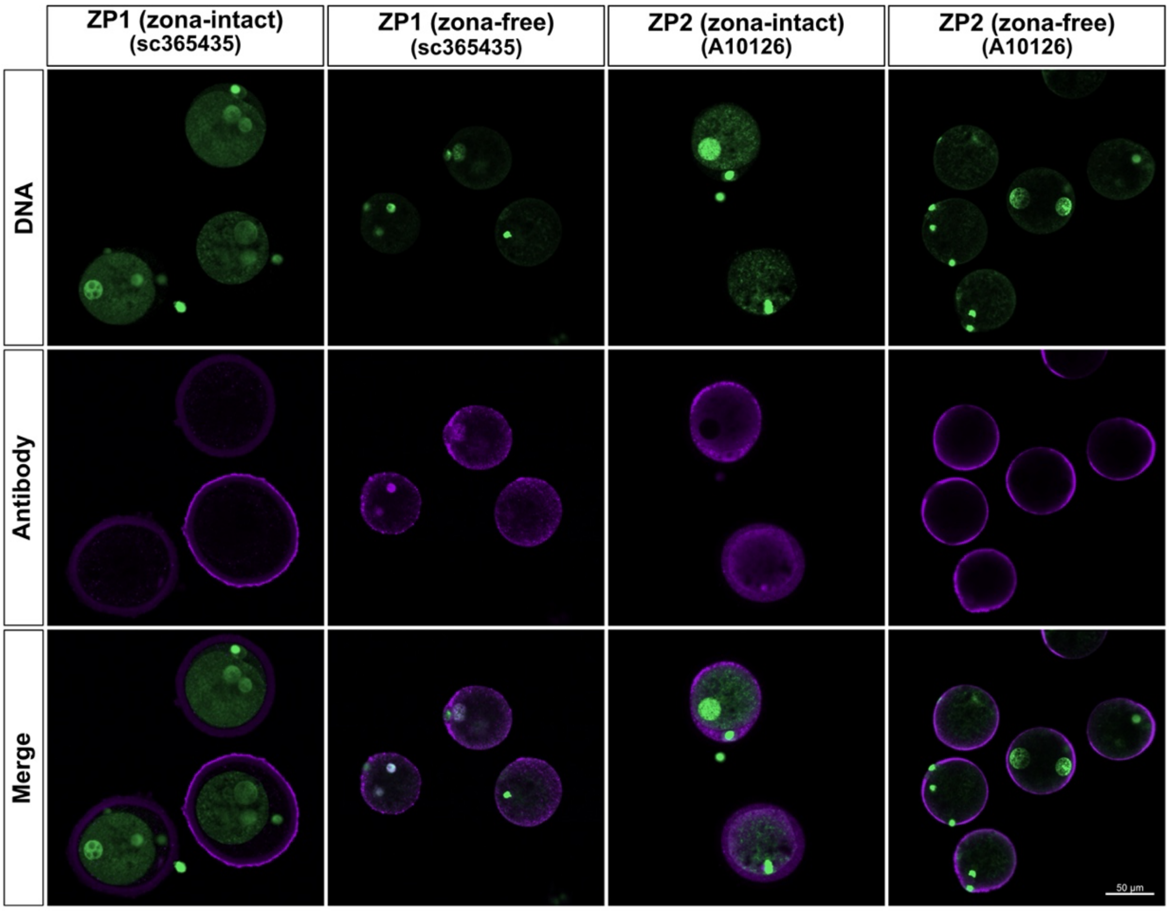


**Supplementary Figure S3. Immunofluorescent stainings of zona-intact and zona-free embryos of various preimplantation stages, revealing that the intracellular ZP signal is not limited to ZP3 but applies also to ZP1 and ZP2. (a).** Immunofluorescence of ZP3 in zona-free embryos. Arrows point at the non-uniform intensity of the peripheral signal. **(b).** Immunofluorescence of ZP1 and ZP2 in zona-free zygotes. Nuclei (DNA) were stained with YO-PRO-1 and are green-fluorescent.

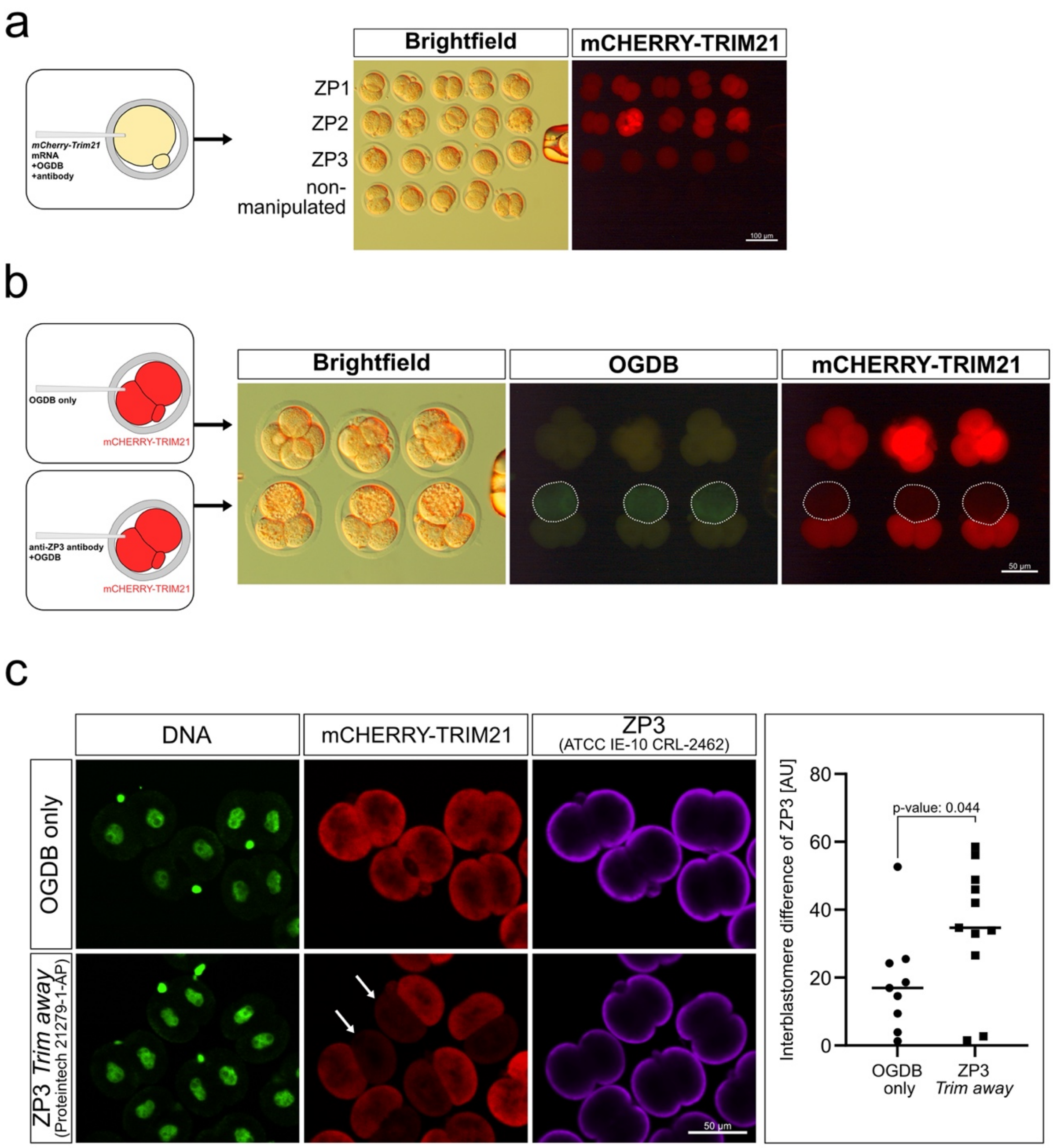


**Supplementary Figure S4. Demonstration of mCHERRY-TRIM21-mediated proteasomal degradation (*Trim-away*) of the ZP proteins, visualized in live (a, b) or fixed (c) embryos using the mCherry tag of TRIM21. (a).** Bright field and fluorescence images of embryos 24 hours after *Trim-away* of ZP1, ZP2 and ZP3 displayed in stacked rows. **(b).** Representative images of embryos preloaded with *mCherry-Trim21* mRNA at the 1-cell stage followed by microinjection of anti-ZP3 in one blastomere (note the drop of mCHERRY fluorescence intensity, dotted circles). **(c).** Demonstration of ZP3 knockdown in 2-cell embryos that were preloaded with *mCherry-Trim21* mRNA at the 1-cell stage and then microinjected with anti-ZP3 (Proteintech 21279-1-AP) in one blastomere only (controls received OGDB in lieu of antibody). *Trim-away* achieved a knockdown. Residual ZP3 was revealed using the antibody ATCC IE-10 CRL-2462, which binds the ZP3 still left after *Trim-away* with Proteintech antibody. Fluorescence intensities were measured in both blastomeres of each embryo, and used to calculate absolute interblastomere differences of ZP3 signal intensity in the *Trim-away vs.* control (OGDB microinjection) groups. Interblastomere differences are higher in the *Trim-away* embryos ($p = 0.044$; Wilcoxon test). The raw measurement data are provided Supplementary Table S4. Nuclei (DNA) were stained with YO-PRO-1 and are green fluorescent. AU, arbitrary units. OGDB, Oregon green dextran beads, co-injected as tracer. mCherry, fluorescent tag of TRIM21.

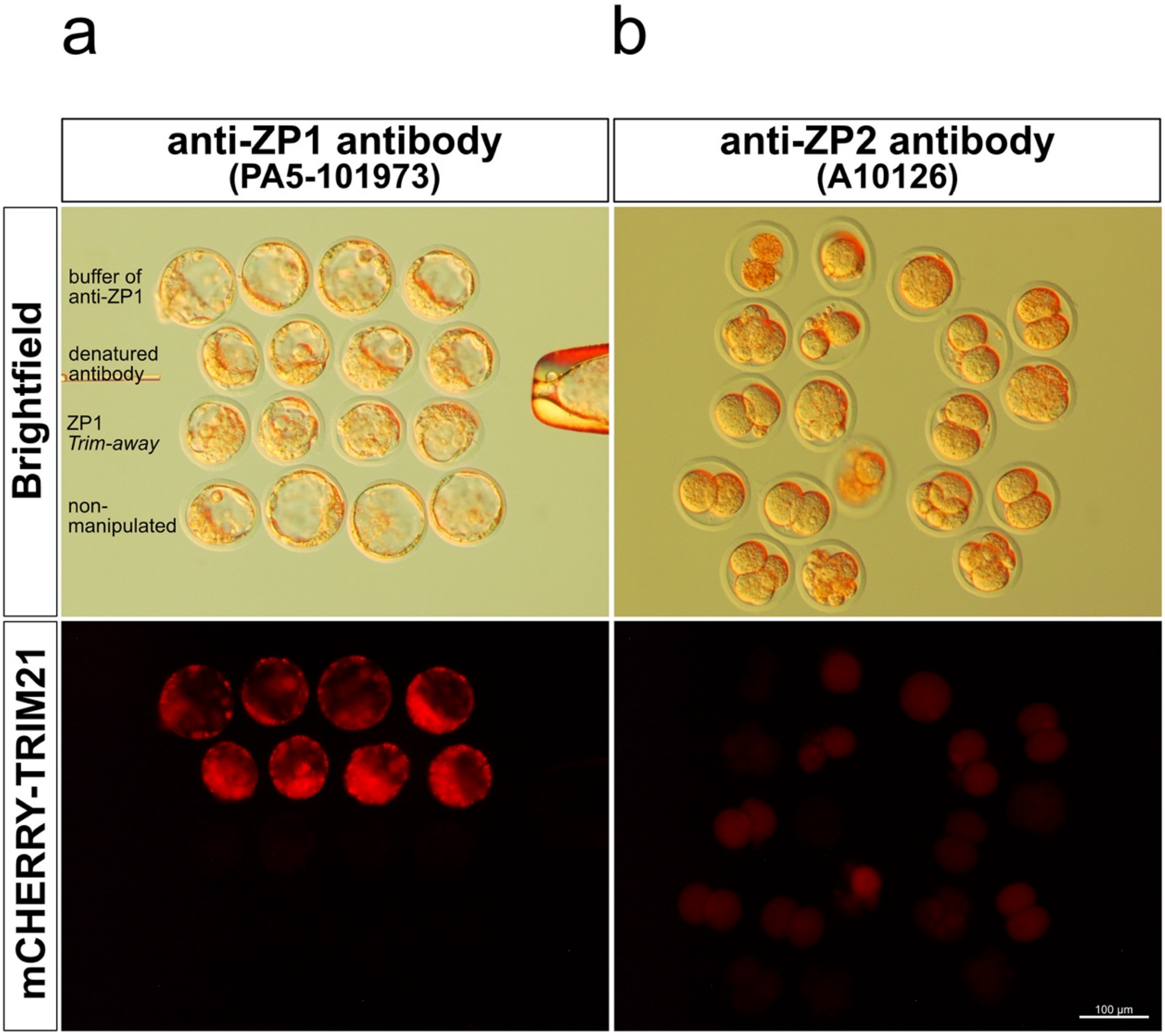


**Supplementary Figure S5. Embryopathy observed after *Trim-away* of the ZP1 and ZP2 proteins. (a).** Stunted blastocyst progression for ZP1 in contrast to **(b).** earlier arrest at the 2-cell or 4-cell stage for ZP2. Note the outwardly normal blastocyst formation **(a).** after microinjection of the heat-denatured antibody or the antibody obtained with the 3$^{rd}$ flow of the Amicon filter devices used to purify the antibody. mCherry, fluorescent tag of TRIM21.

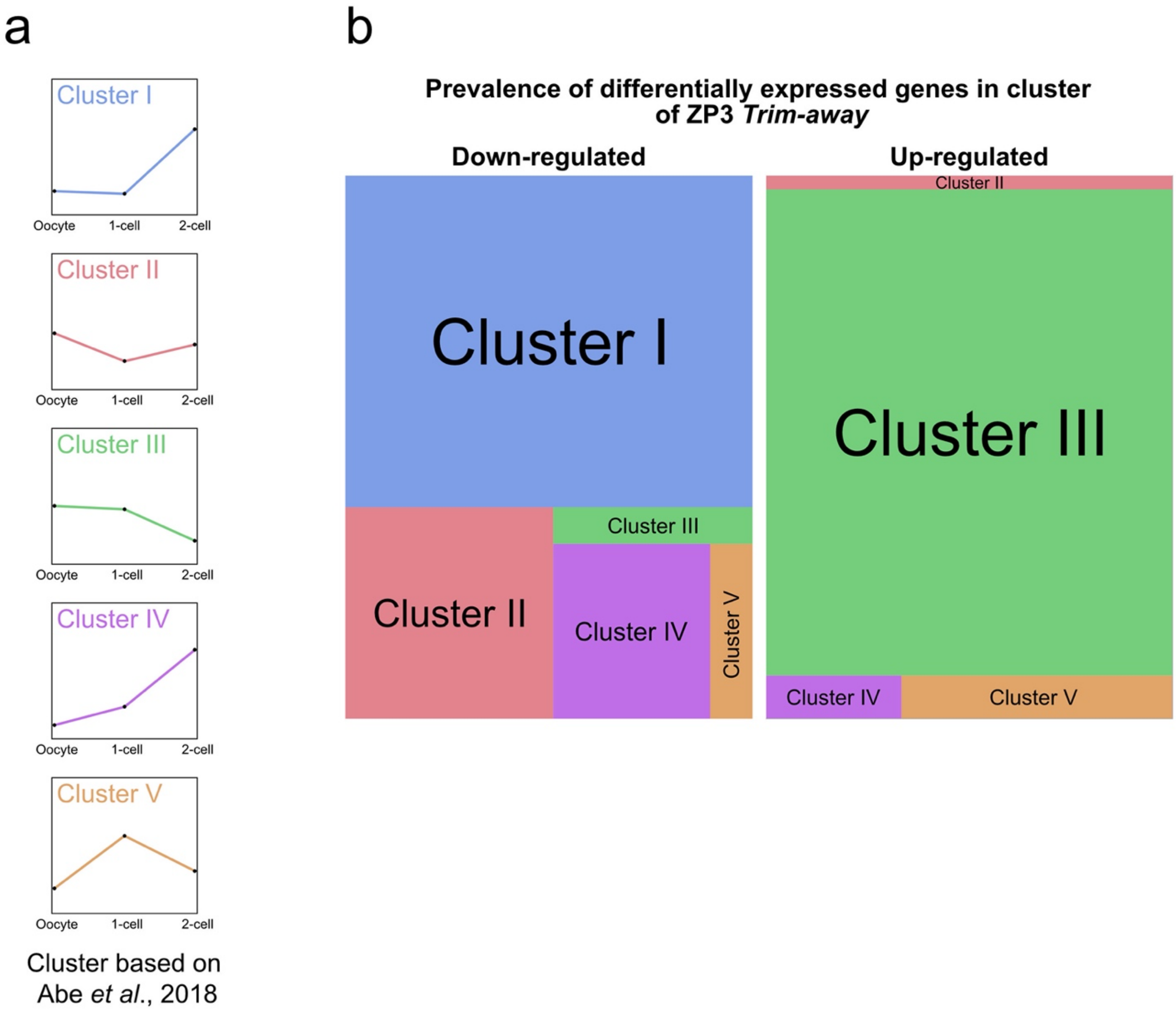


**Supplementary Figure S6. Allocation of the differentially expressed mRNAs of ZP3-knockdown to predefined EGA gene clusters. (a).** Clusters of EGA-regulated genes as defined by Abe *et al.* (2018) according to the increasing or decreasing mRNA abundance during the 1- and 2-cell stage. **(b).** Tree maps of the mRNAs differentially expressed after ZP3-knockdown, showing how they relate to the EGA clusters of Abe *et al.* (2018). 'Down-regulated' means that the mRNAs differentially expressed following ZP3-*Trim-away* are less expressed while in normal embryos they are up-regulated (cluster I). 'Up-regulated' means that the differentially expressed mRNAs are higher expressed while in normal embryos they are down-regulated (cluster III).
